# Distinct RhoGEFs activate apical and junctional actomyosin contractility under control of G proteins during epithelial morphogenesis

**DOI:** 10.1101/566919

**Authors:** Alain Garcia De Las Bayonas, Jean-Marc Philippe, Annemarie C. Lellouch, Thomas Lecuit

## Abstract

Small RhoGTPases and Myosin-II direct cell shape changes and movements during tissue morphogenesis. Their activities are tightly regulated in space and time to specify the desired pattern of contractility that supports tissue morphogenesis. This is expected to stem from polarized surface stimuli and from polarized signaling processing inside cells. We examined this general problem in the context of cell intercalation that drives extension of the *Drosophila* ectoderm. In the ectoderm, G protein coupled receptors (GPCRs) and their downstream heterotrimeric G proteins (Gα and Gβγ) activate Rho1 both medial-apically, where it exhibits pulsed dynamics, and at junctions, where its activity is planar polarized (Kerridge et al., 2016; Munjal et al., 2015). However, the mechanisms responsible for polarizing Rho1 activity are unclear. In particular, it is unknown how Rho1 activity is controlled at junctions. We report a division of labor in the mechanisms of Rho1 activation in that distinct guanine exchange factors (GEFs), that serve as activators of Rho1, operate in these distinct cellular compartments. RhoGEF2 acts uniquely to activate medial-apical Rho1. Although RhoGEF2 is recruited both medial-apically and at junctions by Gα_12/13_-GTP, also called Concertina (Cta) in *Drosophila*, its activity is restricted to the medial-apical compartment. Furthermore, we characterize a novel RhoGEF, p114RhoGEF/Wireless (Wrl), and report its requirement for cell intercalation in the extending ectoderm. p114RhoGEF/Wireless activates Rho1 specifically at junctions. Strikingly it is restricted to adherens junctions and is under Gβ13F/Gγ1 control. Gβ13F/Gγ1 activates junctional Rho1 and exerts quantitative control over planar polarization of Rho1. In particular, overexpression of Gβ13F/Gγ1 leads to hyper planar polarization of Rho1 and MyoII. Finally, we found that p114RhoGEF/Wireless is absent in the mesoderm, arguing for a tissue-specific control over junctional Rho1 activity. These results shed light on the mechanisms of polarization of Rho1 activity in different cellular compartments and reveal that distinct GEFs are sensitive tuning parameters of cell contractility in remodeling epithelia.

## Introduction

Contractile actomyosin networks power cell shape changes during tissue morphogenesis (Gorfinkiel and Blanchard, 2011; Martin and Goldstein, 2014; Munjal and Lecuit, 2014). By pulling on actin filaments anchored to E-cadherin complexes at adherens junctions, non-muscle Myosin-II motors (Myo-II) generate tensile forces whose amplitude and orientation determine the nature of cell and tissue level deformation (Collinet and Lecuit, 2013; Heisenberg and Bellaïche, 2013; Kale et al.; Lecuit and Lenne, 2007; Priya and Yap, 2015). Consequently, specific cortical Myo-II patterns predict specific cell shape changes underlying tissue dynamics(Lecuit et al., 2011; Streichan et al., 2018). During *Drosophila* embryogenesis, apical constriction of cells underlies mesoderm invagination (Leptin and Grunewald, 1990; Sweeton et al., 1991). Apical constriction is driven by a strictly medial-apical pool of Myo-II (Martin et al., 2010). In contrast, during elongation of the ventro-lateral ectoderm (also called germ-band extension), cells intercalate as a consequence of a polarized shrinkage of dorso-ventral interfaces or “vertical junctions” (Bertet and Lecuit, 2009; Blankenship et al., 2006; Irvine and Wieschaus, 1994). This process depends on both a medial-apical pulsatile Myo-II pool and a planar-polarized junctional Myo-II pool to remodel cell interfaces during tissue extension (Bertet and Lecuit, 2009; Blankenship et al., 2006; Rauzi et al., 2010).

The small GTPase Rho1 is a chief regulator of actomyosin networks in these developmental contexts (Mason et al., 2013; De Matos Simões et al., 2014; Munjal et al., 2015), though Rac1 can also activate actin in epithelial cells (Sun et al., 2017). Rho1 cycles between an inactive GDP-bound conformation and an active GTP-bound form. Rho1-GTP binds to and thereby activates the kinase Rok which in turn phosphorylates non muscle Myosin-II regulatory light chain (MRLC, *Sqh* in *Drosophila*). This promotes assembly of Myo-II minifilaments on actin filaments and induces contractility of actomyosin networks. Two families of proteins regulate Rho cycling: Rho guanine nucleotide exchange factors (RhoGEFs), which promotes the exchange of GDP to active GTP bound form of Rho1 and Rho GTPase-activating proteins (RhoGAPs) that inactivate Rho1 by promoting GTP hydrolysis to GDP (Cherfils and Zeghouf, 2013). Recent work has explored the contribution of specific GEFs and GAPs during tissue invagination (Greenberg and Hatini, 2011; Mason et al., 2016; Simões et al., 2006). In the mesoderm, apically localized RhoGEF2, the *Drosophila* ortholog of the mammalian RH-RhoGEFs subfamily (p115RhoGEF/PDZ-RhoGEF/LARG) (Aittaleb and Boguth, 2010; Carter et al., 2014; Meyer et al., 2008), and the RhoGAP Cumberland tune and restrict Rho1 signaling to the apical cell cortex (Mason et al., 2016). How Rho1 activity and therefore the Myo-II activity patterns are controlled during cell intercalation where Rho1 is active both medial-apically and at junctions remains unclear.

The Rho1-Rok core pathway activates both medial-apical and junctional Myo-II in the ectoderm (De Matos Simões et al., 2014; Munjal et al., 2015). Activation of Rho1 occurs via different molecular mechanisms in these distinct cellular compartments downstream of G-protein coupled receptors (GPCRs) and their associated heterotrimeric G proteins (Kerridge et al., 2016). Fog, a GPCR ligand initially reported for its function during apical constriction in the mesoderm (Costa et al., 1994; Dawes-Hoang, 2005a; Manning and Rogers, 2014), is also required for cell intercalation in the ectoderm (Kerridge et al., 2016). It is thus a general regulator of medial-apical Rho1 activation in the embryo, mediated by Gα_12/13_/Cta and RhoGEF2. In the *Drosophila* embryo, the Fog-Gα_12/13_/Cta-RhoGEF2 signaling module specifically controls medial-apical Rho1 activity. The secreted Fog ligand binds to GPCRs Smog and Mist whose GEF activity catalyzes the dissociation of active Gα_12/13_/Cta –GTP from Gβγ (Kerridge et al., 2016; Manning et al., 2013). Free Gα_12/13_/Cta-GTP then binds to RhoGEF2, which in turn activates Rho1, Rok and Myo-II at the apical membrane. In the mesoderm, apical targeting of RhoGEF2 activity is driven by both active Gα_12/13_/Cta and enhanced by the mesoderm-specific apical transmembrane protein T48 which binds the PDZ domain of RhoGEF2(Kolsch et al., 2007). Whether Gα_12/13_/Cta is sufficient to localize RhoGEF2 activity medial-apically in the ectoderm, where T48 is not expressed, is unknown.

A separate biochemical module was hypothesized to control and polarize junctional Rho1 independently in the ectoderm but the underlying molecular mechanisms remain unclear. The pair-rule genes *even-skipped* (*eve*) and *runt* were the first upstream regulators of planar polarized junctional Myo-II identified in the ectoderm (Irvine and Wieschaus, 1994; Zallen and Wieschaus, 2004). The Tolls receptors (Toll2/6/8) are transmembrane proteins whose expression in stripes is regulated by Eve and Runt and who are essential for the polarization of Myo-II (Paré et al., 2014). However, the molecular mechanisms linking Tolls to Rho1 activation remain uncharacterized. The GPCR Smog and the two heterotrimeric G protein subunits Gβ13F/Gγ1 are involved in the tuning of Rho1 activity at ectodermal junctions (Kerridge et al., 2016). However, in the absence of a direct junctional Rho1 activator, e.g. a specific RhoGEF, it is difficult to understand how these upstream regulators polarize the GTPase activity. In this study, we aim to dissect the spatial and temporal control of both medial-apical and junctional Rho1 activity in the ectoderm.

## Results

### RhoGEF2 controls medial-apical Rho1 activity in the ectoderm

We used a Rho1-GTP biosensor that consists of a fusion protein between mEGFP (A206K monomeric EGFP) and the Rho binding domain (RBD) of Anillin which binds selectively to active Rho1-GTP ([Ani-RBD::GFP])(Munjal et al., 2015) in the ectoderm. Ani-RBD::GFP localization shows that active Rho1 is present both medial-apically (Fig.1a, top panel right) and at adherens junctions (Fig.1a, bottom panel left) where it is planar polarized (white arrowheads) as previously reported (Munjal et al., 2015). Importantly, the Rho1 activity pattern is not a consequence of a differential subcellular enrichment in Rho1 protein. Indeed, Rho1 is uniformly distributed along cell membrane in contrast to the planar polarized Rho1-GTP biosensor (Fig. S1 a-c). Hence, Rho1 regulators spatially control Rho1 activity in this tissue. RhoGEF2 is a major activator of the medial-apical Myo-II pool, but not the junctional pool in the ectoderm (Kerridge et al., 2016). Therefore, we first asked whether medial-apical Rho1 activity is specifically decreased upon *RhoGEF2* knock-down. The Rho1-GTP biosensor was analyzed in embryos expressing shRNA against RhoGEF2 driven by maternally supplied Gal4 (matα-Gal-VP16). We found that Rho1-GTP was indeed decreased apically but strikingly preserved at junctions (Fig. 1, b-d, Supplementary Movie 1), consistent with the specific regulation of medial-apical MyoII by RhoGEF2 previously described (Kerridge et al., 2016).

**Figure 1.**
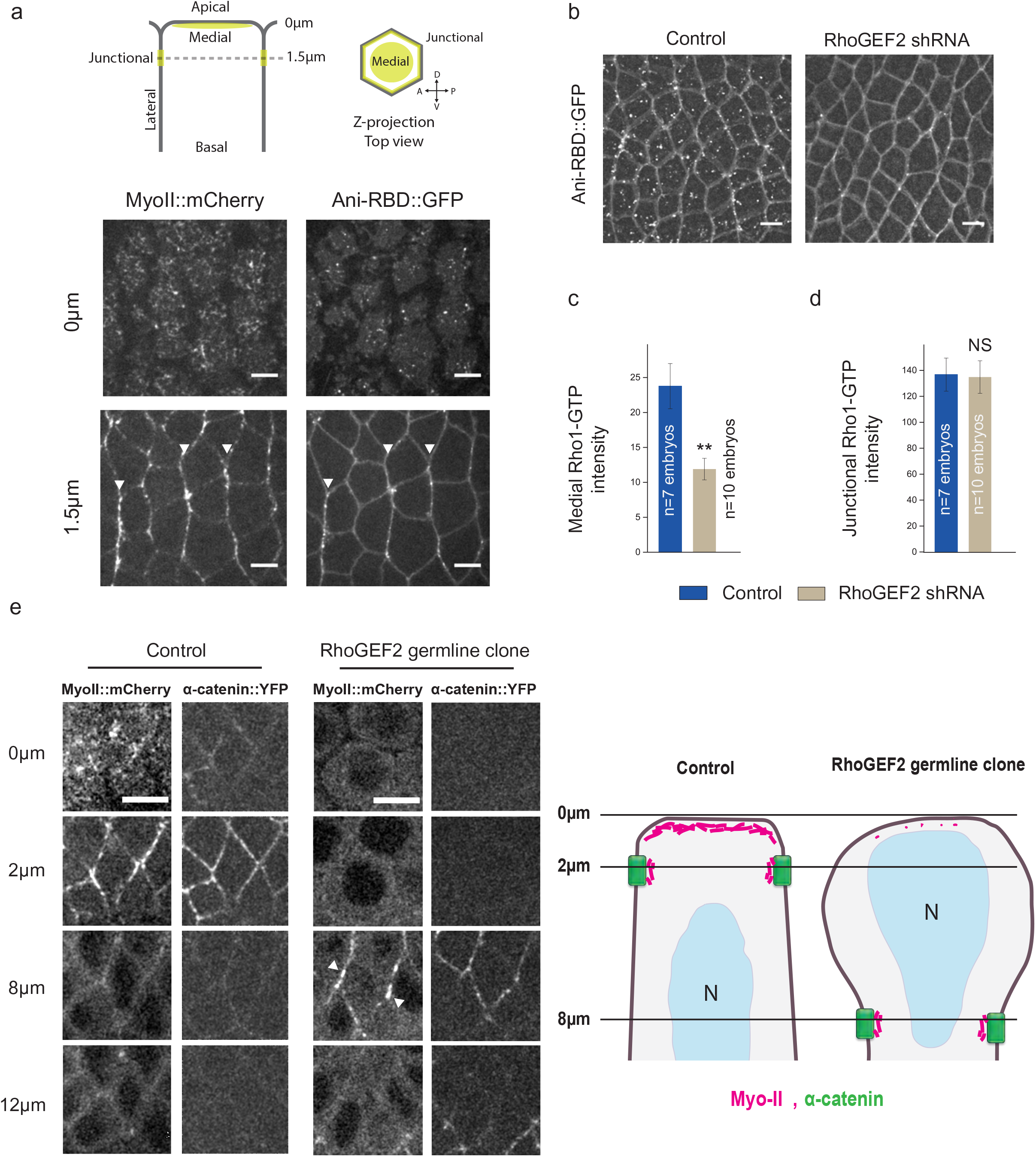
RhoGEF2 activates medial-apical but not junctional Rho1 in the ectoderm. (a) Apical (0µm) and junctional (1.5µm) confocal z-sections of ventro-lateral ectodermal cells from embryos expressing MyoII::mCherry and Ani-RBD::GFP, 8min after the onset of cephalic furrow formation. White arrowheads show planar-polarized Myo-II and Rho1-GTP at vertical junctions. (b) 7µm projections of confocal acquisitions in both control and RhoGEF2 shRNA embryos expressing Ani-RBD::GFP. (c,d) Quantifications of mean medial-apical Rho1-GTP and mean junctional Rho1-GTP intensities in control and RhoGEF2 shRNA embryos. (e) Left panels: Apical (0µm), junctional (2µm) and lateral (8 and 12µm) confocal z-sections of ectodermal cells in control and RhoGEF2 germline clone embryos expressing MyoII::mCherry and α-catenin::YFP, a junctional marker. Medial-apical Myo-II is lost in mutant embryos while Myo-II is still detected at junctions in this condition (white arrowheads). Although half of the RhoGEF2 germline clone embryos express RhoGEF2 zygotically, no rescue has been observed for Myo-II apical levels suggesting that maternally-loaded RhoGEF2 mainly controls the process in a wild-type embryo at this stage. Right panel: schematic view of Myosin-II and adherens junction distribution in both control and RhoGEF2 mutant ectodermal cells. Scale bars = 5µm. Means ± SEM between images are shown. Statistical significance has been calculated using Mann-Whitney U test. ns, p>0.05; * p<0.05; ** p<0.01. All the panels have the same orientation: dorsal at the top, anterior to the left.

These shRNA studies could not rule out a residual RhoGEF2 population signaling at junctions. Therefore, we generated RhoGEF2 maternal and zygotic mutants with germline clones using a null allele for *RhoGEF2*, *DRhoGEF2^l(2)04291^* (Häcker and Perrimon, 1998), and observed a complete loss of medial-apical Myo-II together with an expanded cell surface area (Fig. 1, e). Interestingly, junctional Myo-II persisted in RhoGEF2 mutant embryos. Adherens junctions were found deeper in the tissue relative to wild type junctions, consistent with a role of apical contractility in the positioning of apical junctions (Dawes-Hoang, 2005b; Weng and Wieschaus, 2016). Thus, loss of RhoGEF2 affects medial-apical but not junctional Rho1 signaling. Overall, in the ectoderm, RhoGEF2 is specifically required for Rho1 medial-apical activation, but not for junctional activation.

### Regulation of RhoGEF2 localization and activity in the ectoderm

The spatial distribution of Rho1 signaling could stem from specific control over the localization and/or activity of upstream Rho1 regulators (Mason et al., 2016; Simões et al., 2006). Therefore, we analyzed RhoGEF2 localization in the ectoderm by imaging embryos expressing RhoGEF2::GFP (Mason et al., 2016), whose expression rescues early embryonic phenotypes in RhoGEF2 mutants, and Myo-II::mCherry. RhoGEF2 was enriched both apically and at cell junctions (Fig. 2a), in agreement with previous reports (Levayer et al., 2011; Mason et al., 2016). Additionally, we detected a highly dynamic pool of RhoGEF2 « comets » in the cytoplasm (Fig. 2 a, middle right panel, yellow arrowheads) consistent with the observation that RhoGEF2 localizes at microtubule growing (plus) ends in S2 culture cells (Rogers et al., 2004). To test this further i*n vivo*, we analyzed embryos co-expressing RhoGEF2::RFP and GFP-tagged EB1, a microtubule plus end tracking protein, and found that indeed RhoGEF2::RFP co-localizes with EB1::GFP comets (Fig. 2b, Supplementary Movie 2). The much broader spatial distribution of RhoGEF2 with respect to where RhoGEF2 is specifically required for Rho1 activation led us to ask whether RhoGEF2 activity is spatially segregated in the ectoderm.

**Figure 2.**
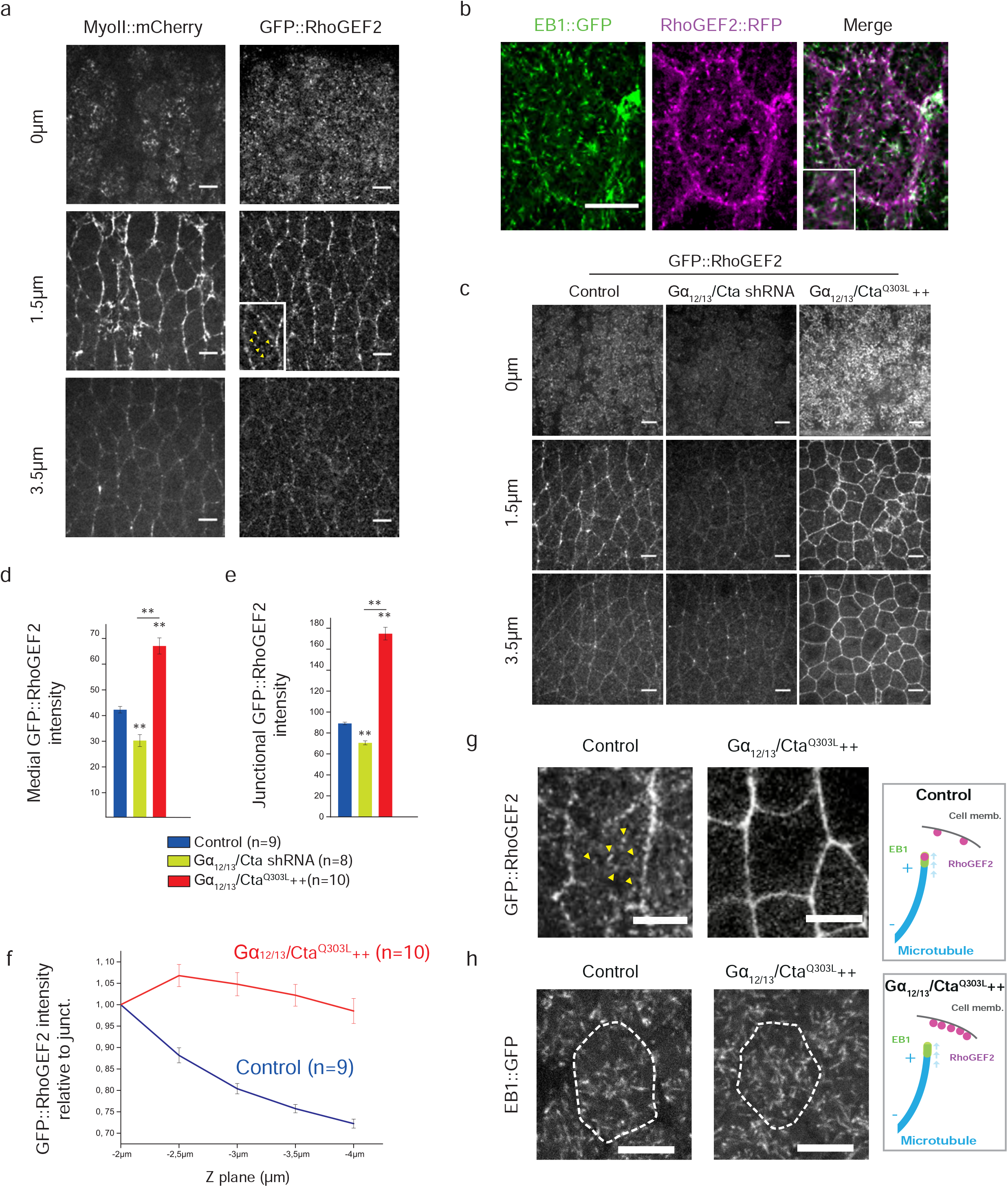
Gα_12/13_/Cta-GTP and microtubules control RhoGEF2 enrichment at cell membrane in the ectoderm. (a) Apical (0µm), junctional (1.5µm) and lateral (3.5µm) z-sections of ectoderm tissue co-expressing MyoII::mCherry and GFP::RhoGEF2. Middle right panel, bottom left corner: a closeup of a cell showing GFP::RhoGEF2 « comets » (yellow arrowheads) (b) Confocal Z-section of an ectodermal cell co-expressing RhoGEF2::RFP and EB1::GFP. Both EB1 and RhoGEF2 « comets » colocalize. (c) Apical (0µm), junctional (1.5µm) and lateral (3.5µm) z-sections of ectoderm tissue expressing GFP-RhoGEF2 in control, Gα_12/13_/Cta shRNA and Gα_12/13_/Cta^Q303L^++ embryos. (d and e) Quantifications of mean medial-apical and junctional GFP::RhoGEF2 intensities in control, Gα_12/13_/Cta shRNA and Gα_12/13_/Cta ^Q303L^++ embryos. n=number of embryos (f) Total GFP::RhoGEF2 cortical levels normalized to the apical junctional intensities (−2µm below the apical membrane). (g) Confocal Z-section of an ectodermal cell expressing GFP::RhoGEF2 in control and Gα_12/_13/Cta^Q303L^++ embryos. GFP::RhoGEF2 « comets » (yellow arrowheads) are absent from Gα_12/_13/Cta^Q303L^++ embryos. (h) Confocal cross-section of an ectodermal cell expressing EB1::GFP in control and Gα_12/_13/Cta^Q303L^++ embryos. Scale bars = 5µm. Means ± SEM between images are shown. Statistical significance has been calculated using Mann-Whitney U test. ns, p>0.05; * p<0.05; ** p<0.01. All the panels have the same orientation: dorsal at the top, anterior to the left.

Gα_12/13_/Cta and the membrane anchor T48 promote RhoGEF2 activation at the cell membrane in *Drosophila* upon GPCR activation (Kölsch et al., 2007; Rogers et al., 2004). Both regulators cooperate to recruit RhoGEF2 to the apical membrane in the mesoderm where it activates Rho1 signaling. T48 anchors RhoGEF2 via a direct PDZ domain interaction. By analogy to its mammalian homolog p115RhoGEF, RhoGEF2 is thought to bind to active Gα_12/13_/Cta via its N-terminal RH domain. A conformational change then dislodges the autoinhibitory N-terminal tail of the RhoGEF from its DH-PH domains, making them accessible for binding to Rho1 and membrane lipids (Aittaleb and Boguth, 2010). Although this allosteric regulation by active Gα_12/13_/Cta is sufficient to increase p115RhoGEF binding to the membrane, it is not clear whether a full activation of the RhoGEF requires additional control. T48 is not expressed in the ectoderm (Kölsch et al., 2007) and therefore cannot account for RhoGEF2 activity at the apical membrane though T48 overexpression in the ectoderm can increase apical Myo-II activation (data not shown) similar to RhoGEF2 overexpression (Kerridge et al., 2016). We previously showed that RhoGEF2 is epistatic to Gα_12/13_/Cta in the extending lateral ectoderm (Kerridge et al., 2016). Indeed, the Gα_12/13_/Cta_-_dependent increase of medial-apical Myo-II is abolished upon *RhoGEF2* knock-down, indicating that RhoGEF2 transduces the signal downstream of Gα_12/13_/Cta. Therefore, Gα_12/13_/Cta is a strong candidate for controlling RhoGEF2 localization and activity in the ectoderm. We examined RhoGEF2 localization in Gα_12/13_/Cta-depleted embryos and in embryos expressing constitutively active Gα_12/13_/Cta, Gα_12/13_/Cta ^Q303L^ (a mutant that mimics the GTP bound state). Apical and junctional RhoGEF2 levels in Gα_12/13_/Cta knock-down embryos (Fig. 2c) significantly decreased (Fig.2 d and e). This shows that Gα_12/13_/Cta is required to localize RhoGEF2 in both compartments. Strikingly in Gα_12/13_/ Cta ^Q303L^ embryos, RhoGEF2 was strongly enriched everywhere at the cell surface, namely the apical membrane, at junctions and along the lateral cell surface (Fig. 2c-e). In contrast, RhoGEF2 « comets » were completely absent from the cytoplasm in this condition (Fig. 2 g, yellow arrowheads and Supplementary Movie 3) while EB1 comets were still present as in controls (Fig. 2 h). This suggests that Gα_12/13_/Cta-GTP promotes RhoGEF2 dissociation from microtubule growing ends and its subsequent enrichment at cell membrane upon GPCR activation, as reported in S2 cells (Rogers et al., 2004).We further tested whether microtubules sequester RhoGEF2 and thereby limit RhoGEF2 membrane recruitment and signalling. Microtubule depolymerization following injection of colcemid caused germ-band extension defects (Fig. S2 a and b) and a medial-apical increase in Myo-II activation (Fig. S2 c and d). This phenotype was similar to RhoGEF2 or Gα_12/13_/Cta overexpression (Kerridge et al., 2016), arguing that microtubules sequester and thereby limit RhoGEF2 signaling medial-apically. Note that while medial-apical Rho1-GTP levels increased in Gα_12/13_/ Cta ^Q303L^ expressing embryos, they were unchanged at junctions (Fig. S2 e-g), consistent with the previous report showing that only medial-apical Myo-II was affected in such conditions (Kerridge et al., 2016). Thus, although active Gα_12/13_/Cta releases RhoGEF2 from microtubule plus ends and recruits it both medial-apically and at junctions in the wild type and in over-expression conditions, RhoGEF2 signaling is consistently restricted to the apical membrane.

### Identification of a new RhoGEF required during tissue extension

The striking apical specificity of RhoGEF2 indicates that other RhoGEF(s) activate junctional Rho1 in the ectoderm. We screened all 26 predicted *Drosophila* RhoGEFs for defects in germ-band extension by expressing shRNA maternally and zygotically. Knock-down of the maternal contribution was crucial in such experiments, as a strong maternal mRNA loading is observed for a large number of RhoGEFs in the embryo (modENCODE_mRNA-Seq, Flybase). Knock-down of *CG10188* slowed germ-band extension (Fig. 3 a and b). Notably, intercalation events, (also called T1 events), which underlie tissue extension (Claire Bertet, 2004; Collinet et al., 2015) were significantly decreased in *CG10188* shRNA expressing embryos (Fig. 3 c and d, Supplementary Movie 3). Severe developmental defects were also observed at later stages such as the absence of germ-band retraction and the occurrence of cell delamination resulting in a fully penetrant embryonic lethality (data not shown). We designed a transgene that ubiquitously expresses a modified form of the CG10188 mRNA immune to targeting by the shRNA athough with preserved codon usage (SqhPa-CG10188-shRNA^R^, see Material and Methods). This transgene rescued lethality in *CG10188* shRNA expressing embryos and proved the specificity of the knock-down (Fig. 3 f). Overall, these results demonstrate a requirement for CG10188 during germ-band extension.

**Figure 3.**
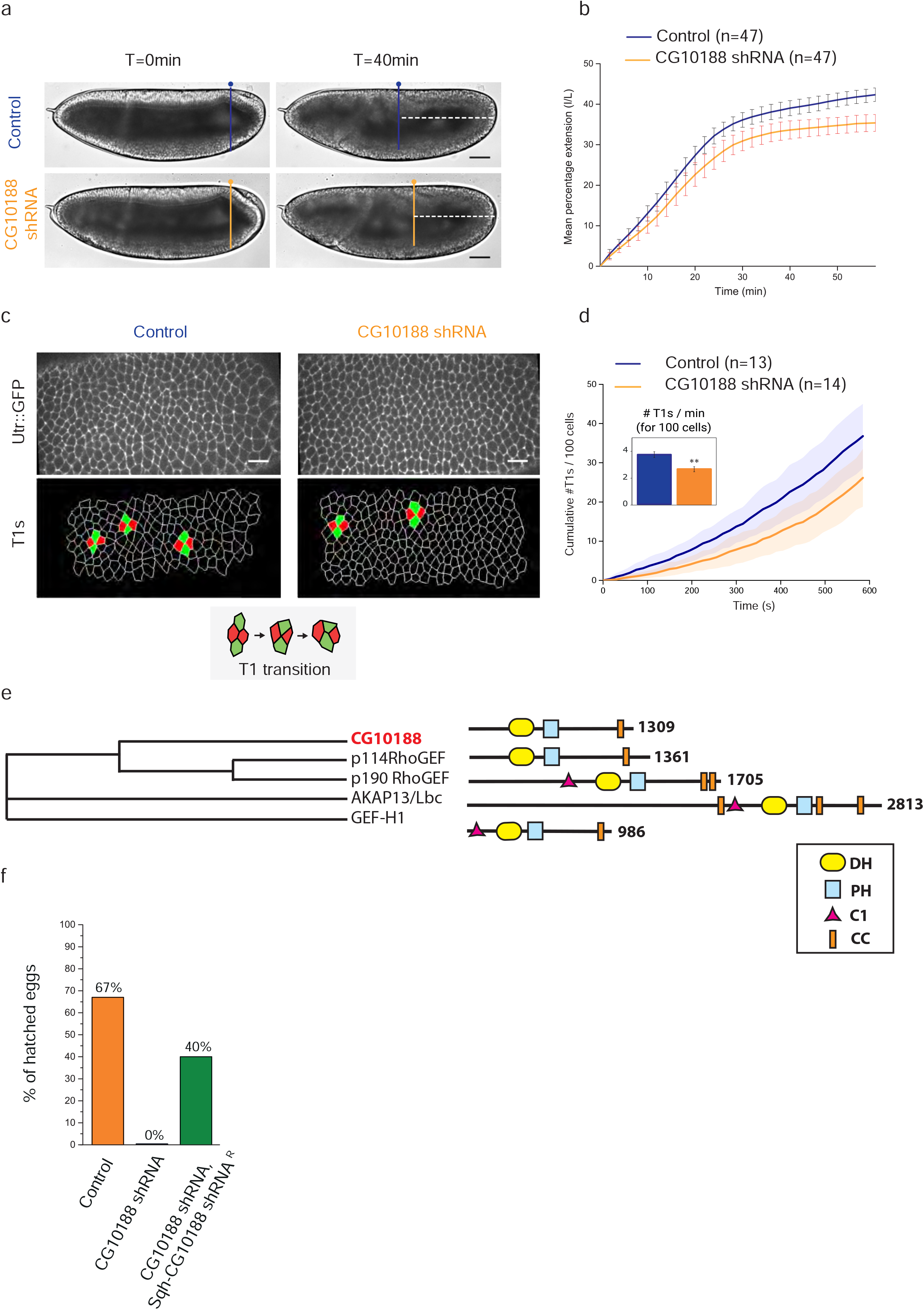
A new RhoGEF controls cell intercalation during germ-band extension. (a) Lateral view of a control and a CG10188 shRNA expressing embryo at the onset (t=0min) of germ-band extension (GBE) and 40min later. The dotted lines mark the distance between the pole cells and the posterior side of the embryos 40 minutes after the onset of GBE. (b) Quantification of germ-band extension over time in control and CG10188 shRNA embryos. n=number of embryos. (c) Top panels: a representative view of the extending ventro-lateral ectoderm in control and CG10188 shRNA embryos expressing Utr::GFP. Bottom panels: segmented view of the same embryos. T1 transitions are depicted in green and red on the image. (d) Cumulative sum of T1 transitions measured for control and CG10188 shRNA embryos over a period of 10 minutes. n=number of embryos. (e) Phylogenetic tree inferred from sequence similarity between the *Drosophila* CG10188 and its human orthologs p114RhoGEF, GEF-H1, p190RhoGEF and AKAP13. Human sequences were collected from UniProt and clustered by multiple sequence alignment using ClustalOmega (nj tree, no distance correction). CG10188 exhibits a DH-PH tandem characteristic of the Dbl-RhoGEFs and a Coil-coiled (CC) motif in its C-terminal region, known to be a dimerization domain in its mammalian counterparts.(f) Percentage of embryos that hatched in control (n=238/354 embryos), CG10188 shRNA (n=0/243 embryos) and cg10188 shRNA, Sqh-cg10188^wt^ shRNA (n=86/213 embryos) conditions. The egg-hatching percentage was determined as a measurement of embryo viability (see Materials and Methods). The fully penetrant embryonic lethality observed in cg10188 shRNA embryos is rescued by the expression of the targeted gene refractory to the shRNA (Sqh-CG10188 shRNA^R^). Scale bars = 50µm (a) and =15µm (c). Error bars, SD for b and SEM for d.

CG10188 has not yet been functionally characterized in *Drosophila*. From sequence and domain similarity, CG10188 is the ortholog of the mammalian RhoGEF subfamily including p114RhoGEF, AKAP13, GEF-H1 and p190RhoGEF, who each activate RhoA (Fort and Blangy, 2017; Nakajima and Tanoue, 2011) (Fig.3 e). Based on their sequence and function (Terry et al., 2011) compared with our data hereafter, we conclude that CG10188 is the *Drosophila* functional ortholog of mammalian p114RhoGEF and we will now refer to it as p114RhoGEF. Transcriptomic analyses reported a maternal and zygotic expression of p114RhoGEF in the embryo, suggesting that the protein could be present and active in the ectoderm (Karaiskos et al., 2017; Pilot, 2006).

### p114RhoGEF/Wireless activates Rho1 signaling at adherens junctions in the ectoderm

To test if p114RhoGEF controls Rho1 activity in the ectoderm, we investigated the distribution of the Rho-GTP biosensor in *p114RhoGEF* shRNA expressing embryos. In striking contrast to the RhoGEF2 knock-down, medial-apical Rho1-GTP levels were unaffected whereas junctional Rho1-GTP was strongly decreased (Fig. 4 a-c). The loss of active junctional Rho1 suggested that junctional Myo-II might be affected. Therefore, we analyzed Myo-II::mCherry in control and *p114RhoGEF* shRNA embryos. Similar to Rho1-GTP, junctional Myo-II was strongly reduced while medial-apical Myo-II was preserved (Fig. 4 d-f, Fig. S3 a,b, Supplementary Movie 4). Since planar-polarized Myo-II cables that are associated with the enrichment of Myo-II at vertical junctions were lost in p114RhoGEF shRNA embryos, we named the novel RhoGEF p114RhoGEF Wireless (*wrl*). Interestingly, Myo-II persisted at cell vertices in the *p114RhoGEF/wrl* knock-down (Fig. 4d, bottom right panel). Rho1-GTP is not detected at vertices in this condition (Fig.4 a) which suggests either a redistribution of remaining active Myo-II in this condition or that Myo-II could be activated through different mechanisms in this compartment. Last, compared to wild type, E-cadherin levels were globally reduced in *p114RhoGEF/wrl* knock-down embryos with a highly discontinuous E-cadherin distribution at junctions (Fig. S3 c-e). Similar E-cadherin defects have been observed upon dominant-negative Rho1 expression and Rho1 inhibition (Braga et al., 1999; De Matos Simões et al., 2014; Takaishi et al., 1997), consistent with the specificity of *p114RhoGEF/wrl* for Rho1 signaling.

**Figure 4.**
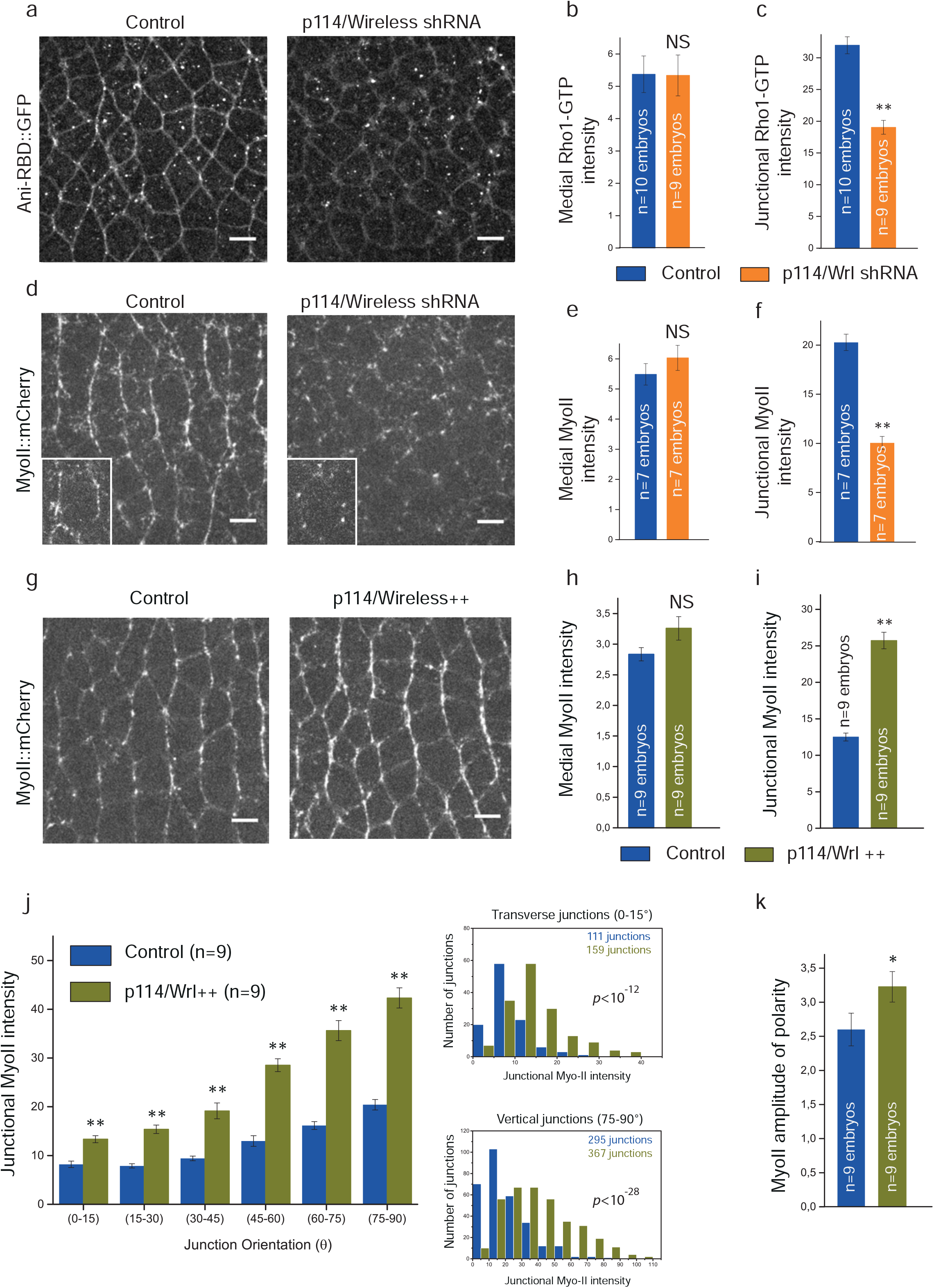
p114RhoGEF/Wireless activates junctional Rho1 signaling in the ectoderm. (a) 4µm z-projection of confocal acquisition of control or p114RhoGEF/Wireless shRNA embryos expressing Ani-RBD::GFP. Active Rho1 is specifically decreased at junctions upon p114RhoGEF/Wireless knock-down. (b, c) Mean medial-apical and junctional Rho1-GTP intensities in both control and p114RhoGEF/Wireless shRNA embryos. (d) Confocal acquisitions of control and p114RhoGEF/Wireless shRNA embryos expressing MyoII::mCherry. A closeup of a representative cell is shown in the bottom part left panel for both conditions (the most apical z-planes containing medial-apical Myo-II have been removed for a better visualization of junctional MyoII::mCherry signal). (e, f) Quantifications of mean medial-apical and junctional Myo-II intensities in both control and p114RhoGEF/Wireless shRNA embryos. (g) MyoII::mCherry in control and Sqh-p114RhoGEF/Wireless embryos (p114RhoGEF/Wireless ++). (h, i) Mean medial-apical and junctional Myo-II intensities in control and p114RhoGEF/Wireless++ embryos. (j) Left panel: Mean junctional intensity of Myo-II according to the angle of the junctions. (junction angle; 0°, parallel to the antero-posterior axis; 90°, perpendicular to the antero-posterior axis). n= number of embryos. Right panels: Distributions of junctional Myo-II intensity values at transverse (0-15°) and vertical junctions (75-90°) in control and p114RhoGEF/Wireless ++ embryos. A significant difference was observed in both angular ranges (Kolmogorov-Smirnov two-sample test). p values for Kolmogorov-Smirnov two-sample test in each comparison are indicated on the plot. ns, not significant, p>0.05. (k) Quantification of Myo-II amplitude of polarity in control and p114RhoGEF/Wireless ++ embryos. Amplitude of polarity is measured as the ratio of mean Myo-II intensity at vertical junctions to mean Myo-II intensity at transverse junctions. Scale bars = 5µm. Means ± SEM are shown. Statistical significance has been calculated using Mann-Whitney U test. ns, p>0.05; * p<0.05; ** p<0.01.

RhoGEF2 exhibits a dose-dependent effect on medial-apical Rho1 signaling in that overexpression of RhoGEF2 is sufficient to increase medial Rho1-GTPase and Myo-II activation(Azevedo et al., 2011; Kerridge et al., 2016). Therefore, we asked whether increasing *p114RhoGEF/wrl* expression levels could, symmetrically, increase Rho1 signaling at junctions. The p114RhoGEF/Wrl levels were increased by driving *p114RhoGEF/wrl* wild type coding sequence under control of the ubiquitous MRLC/*Sqh* promoter in Myo-II::mCherry embryos. The result was unique and striking: p114RhoGEF/Wrl overexpression led to a global Myo-II junctional increase relative to control with no effect on medial-apical Myo-II (Fig. 4 g-i, Supplementary Movie 5). Myo-II was increased both at transverse (0-15°, 63% increase) and vertical junctions (75-90°, 200% increase) (Fig.4 j), with a resulting modest (24%) increase in planar polarity (Fig.4 k). Thus, p114RhoGEF/Wrl tunes Rho1 signaling in a dose dependent manner at junctions.

RhoGEF2 and p114RhoGEF/Wrl show complementary spatial restriction of activity on Rho1 signaling. We thus hypothesized that a double knock-down of both RhoGEFs should abolish total Rho1 activity in the ectoderm. Indeed, Rho1-GTP and Myo-II were decreased both apically and at cell junctions in this context (Fig.S4 a-f). Together, our data demonstrate that p114RhoGEF/Wrl is a key activator of Rho1 signaling at adherens junctions in the ectoderm. Moreover, RhoGEF2 and p114RhoGEF/Wrl have additive and non-redundant functions in the ectoderm.

### p114RhoGEF/Wrl mediates Gβ13F/Gγ1-dependent junctional Rho1 signaling

Given the critical function of Gβ13F/Gγ1 in the regulation of medial-apical and junctional Myo-II pools (Kerridge et al., 2016), we examined its link with p114RhoGEF/Wireless at junctions. We first tested whether Rho activity was dependent upon Gβ13F/Gγ1. We analyzed the Rho1-GTP biosensor distribution in both Gβ13F/Gγ1 loss of function (Gγ1 germline clone) and gain of function (Gβ13F/Gγ1 overexpression) conditions. Loss of Gγ1 resulted in a reduction of both junctional and medial-apical Rho1-GTP, consistent with the overall reduction in Myo-II previously reported (Kerridge et al., 2016) (Fig. 5, a-c). Note that the medial-apical decrease in Rho1 signaling does not imply direct Gβ13F/Gγ1 activity apically as this is expected from the known mechanisms controlling heterotrimeric G protein activation. Indeed the Gβγ subunit dimer is necessary to properly localize Gα at the membrane and thereby to prime Gα to respond to GPCR GEF activity (Evanko et al., 2000; Fishburn et al., 2000; Tang et al., 2006). Thus, Gβ13F/Gγ1 is required for Gα_12/13_/Cta activation (Gα-GTP) downstream of GPCRs such that loss of Gβ13F/Gγ1 also causes loss of Gα_12/13_/Cta activity.

**Figure 5.**
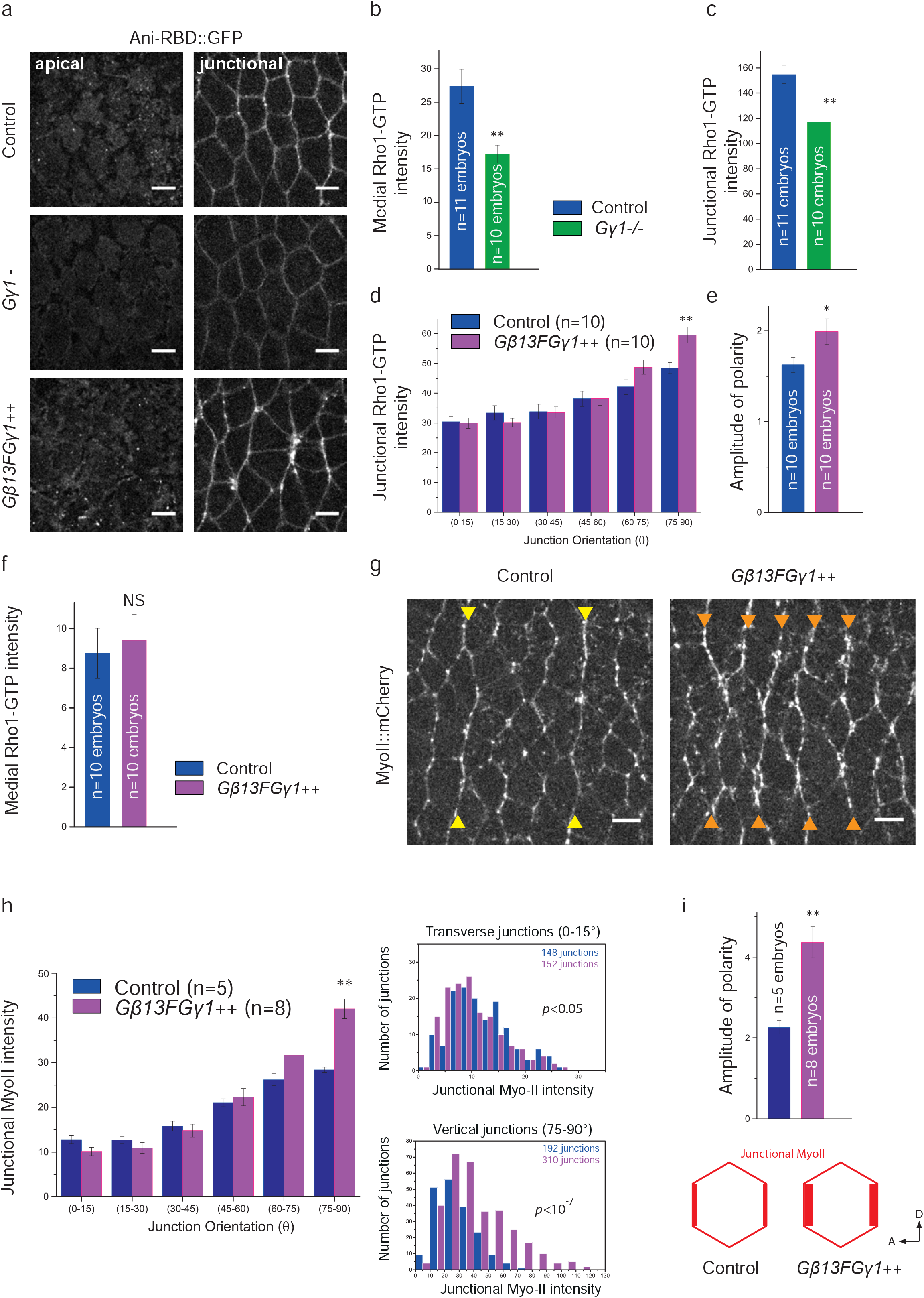
Gβ13FGγ1 activates and polarizes junctional Rho1 signaling in the ectoderm. (a) Apical (0µm) and junctional (1.5µm) confocal z-sections of ventro-lateral ectodermal cells expressing Ani-RBD::GFP in control, Gγ1 germline clone (Gγ1^-^) and Gβ13FGγ1 overexpressing embryos (Gβ13FGγ1++). (b, c) Mean medial-apical and junctional Rho1-GTP intensities in control and Gγ1^-^ embryos. (d) Mean junctional intensity of Rho1-GTP according to the angle of the junctions in control and Gβ13FGγ1++ embryos. n= number of embryos. (e) Rho1-GTP amplitude of polarity. (f) Medial-apical Rho1-GTP intensities in control and Gβ13FGγ1++ embryos. (g) Confocal acquisitions (4.5µm projections) showing MyoII::mCherry in control and Gβ13FGγ1++ embryos. (yellow arrowheads show two strong compartment boundaries cables of Myo-II in a control embryo; orange arrowheads show the ectopic supracellular cables of Myo-II induced by Gβ13FGγ1 overexpression. (h) Left panel: Mean junctional intensity of MyoII::mCherry according to the angle of the junctions in control and Gβ13FGγ1++ embryos. n= number of embryos. Right panels show the distributions of junctional Myo-II intensity values for transverse (0-15°) and vertical junctions (75-90°) in control and Gβ13FGγ1++ embryos. We observed a mild statistical difference at the transverse junctions (Kolmogorov-Smirnov two-sample test, *p*<0.05). This is explained by an increase in the lower Myo-II intensity values in Gβ13F/Gγ1++embryos as compared to control. At vertical junctions, a strong statistical difference was observed (*p*<10^−7^) as a consequence of a global increase in Myo-II intensity values in Gβ13F/Gγ1++embryos compared to control. p values for Kolmogorov-Smirnov test in each comparison are indicated on the plot. ns, not significant, p>0.05. (i) Quantification of Myo-II amplitude of polarity in control and Gβ13FGγ1++ embryos. Scale bars = 5µm. Means ± SEM are shown. Statistical significance has been calculated using Mann-Whitney U test. ns, p>0.05; * p<0.05; ** p<0.01.

We then overexpressed both Gβ13F and Gγ1 in embryos to test a dose-dependent effect of these subunits on junctional Rho1 signaling. Overexpression of either Gβ13F or Gγ1 alone did not give any phenotype (data not shown), consistent with studies showing that the individual Gβ and Gγ subunits can neither be transported to the membrane individually nor bind to or signal via their molecular effectors as monomers (Lukov et al., 2005; Smrcka, 2008). In contrast and remarkably, Gβ13F/Gγ1 co-expression resulted in a specific enrichment in Rho1 activity at vertical junctions (23% increase) compared to controls (Fig.5 d). Consequently, Rho1-GTP planar-polarity was significantly increased (25% increase, Fig.5 e). However, medial-apical Rho1 activity was not significantly changed upon Gβ13F/Gγ1 co-expression, indicating a different sensitivity to Gβ13F/Gγ1 levels in the apical compared to the junctional compartments (Fig.5 f). Note that, Gα_12/13_/Cta showed the opposite pattern (Fig. S2, e-g). Myo-II::mCherry was next examined in Gβ13F/Gγ1 overexpressing embryos (referred to as Gβ13F/Gγ1++). Consistent with the previous data, we observed a specific increase of Myo-II at vertical junctions (48% increase, Fig. 5, g and h, Supplementary Movie 6) leading to a strong (two-fold) increase in Myo-II planar polarity (Fig. 5, i). Because Gβ13F/Gγ1 overexpression hyperpolarized Myo-II in all the ectodermal cells, the strong parasegmental boundaries cables (Tetley et al., 2016) observed in the wild-type (yellow arrowheads in Fig.5 g, left panel) were now indistinguishable from the other vertical interfaces in this condition (orange arrowheads Fig.5 g, right panel). Altogether, we uncovered a new role for Gβ13F/Gγ1 dimer which is involved quantitatively in the planar-polarization of Rho1 signaling at junctions. Therefore, both Gβ13F/Gγ1 and p114RhoGEF/Wrl regulate junctional Myo-II by quantitatively tuning Rho1 activation at junctions.

These results suggested that p114RhoGEF/Wrl might be genetically epistatic to Gβ13F/Gγ1. Thus, we investigated Gβ13F/Gγ1 overexpression in conjunction with *p114RhoGEF/wrl* shRNA to explore this relationship. To avoid any differential titration of Gal4 effects, the number of UAS regulatory sequences was equivalent in both the Gβ13F/Gγ1++ and the Gβ13F/Gγ1++, p114RhoGEF/Wrl shRNA embryos (see Table 1). The polarized increase in Myo-II at vertical junctions in Gβ13F/Gγ1++ embryos was no longer observed in Gβ13F/Gγ1++, p114RhoGEF/Wrl shRNA embryos (Fig.6a) which were indistinguishable from p114RhoGEF/Wrl shRNA embryos alone (Fig.6 b, c compare with Fig.4 d and Fig.S3 a,b). Overall, these data show that p114RhoGEF/Wrl is crucial to mediate Gβ13F/Gγ1-dependent Rho1 signaling at junctions.

**Figure 6.**
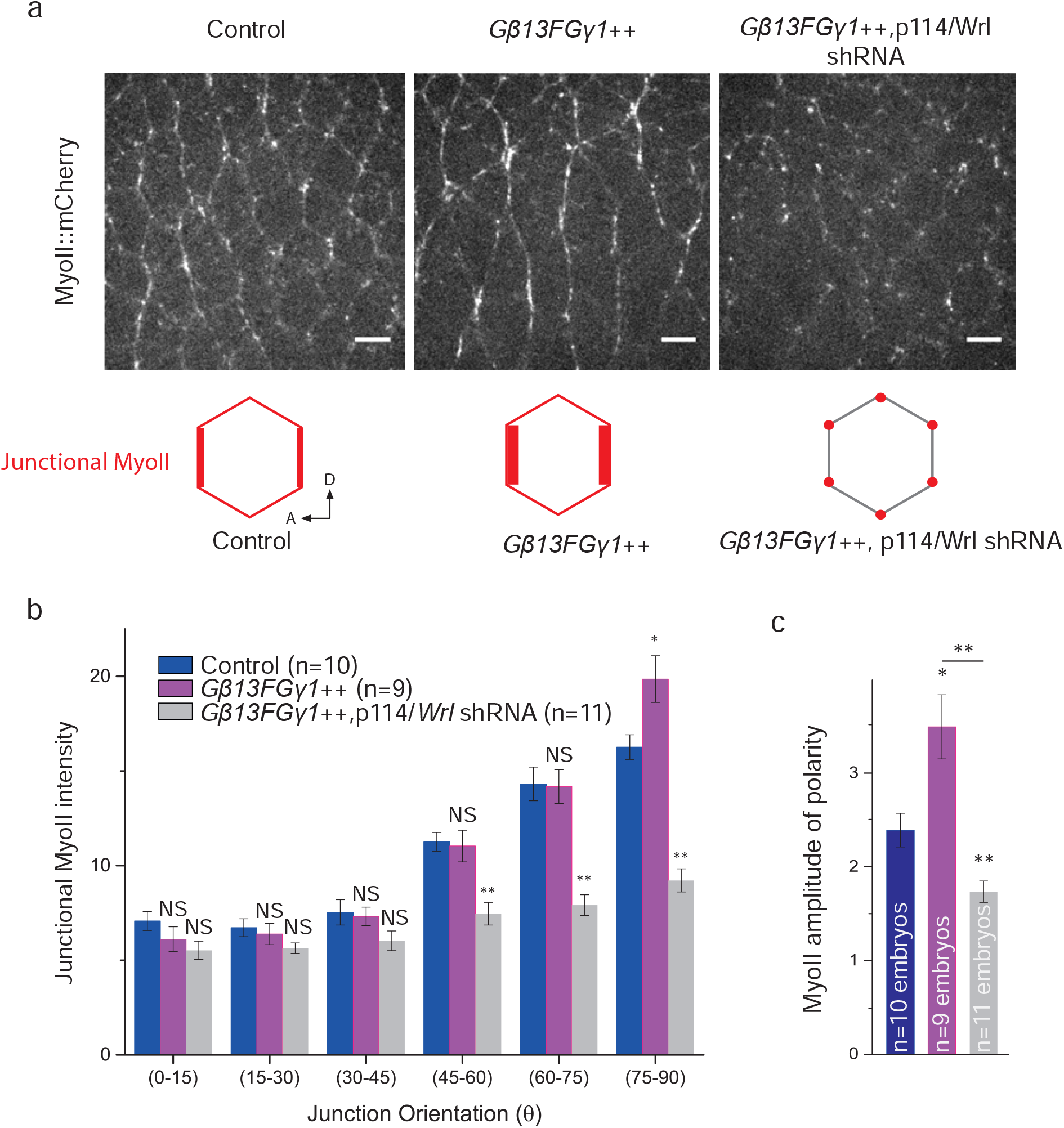
p114RhoGEF/Wireless mediates Gβ13FGγ1 signaling at junctions. (a) Confocal acquisitions of MyoII::mCherry in the ventro-lateral ectoderm of control, Gβ13FGγ1++ and Gβ13FGγ1++, p114/Wrl shRNA embryos. The increase of Myo-II at vertical junctions observed in Gβ13FGγ1++ embryos is lost when p114/Wrl shRNA is also expressed in the background. (b) Mean junctional intensity of MyoII::mCherry according to the angle of the junctions in control, Gβ13FGγ1++ embryos and Gβ13FGγ1++, p114/Wrl shRNA embryos. n= number of embryos. (c) Amplitude of polarity of junctional MyoII::mCherry in control, Gβ13FGγ1++ embryos and Gβ13FGγ1++, p114/Wrl shRNA embryos. While Myo-II planar polarity increases upon Gβ13FGγ1 overexpression compared to control embryos, co-expression of Gβ13FGγ1 together with p114/Wireless shRNA reduces Myo-II planar polarity, similar to p114/Wireless shRNA embryos alone. Scale bars = 5µm. Means ± SEM are shown. Statistical significance has been calculated using Mann-Whitney U test. ns, p>0.05; * p<0.05; ** p<0.01.

### Gβ13F/Gγ1 regulate p114RhoGEF/Wrl junctional enrichment in the ectoderm

The new genetic interaction between Gβ13F/Gγ1 and p114RhoGEF/Wrl led us to ask whether Gβ13F/Gγ1 subunits could activate and/or localize p114RhoGEF/Wrl at junctions. First, we assessed their respective subcellular distribution *in vivo*. Transgenic lines that express p114RhoGEF/Wrl tagged with either N-terminal or C-terminal GFP were generated (see Material and Methods). Embryos expressing GFP-tagged p114RhoGEF/Wrl and Myo-II::mCherry were imaged. We found that p114RhoGEF/Wrl::GFP localization was restricted to adherens junctions, where it forms puncta, in both N- and C-term GFP fusions (Fig. 7 a, Fig. S5 a). Remarkably, while expressed ubiquitously in the embryo, p114RhoGEF/Wrl::GFP was not detected at junctions in the mesoderm (Fig. 7 b). It has been reported that Rho1 signaling in mesodermal cells is induced medial-apically and absent from junctions (Mason et al., 2016). Therefore, a mesoderm-specific regulation is likely to block junctional Rho1 signaling in this tissue via p114RhoGEF/Wrl mRNA or protein degradation since we failed to detect any increase in cytoplasmic p114RhoGEF/Wrl::GFP signal in these cells (Fig. 7 b).

**Figure 7.**
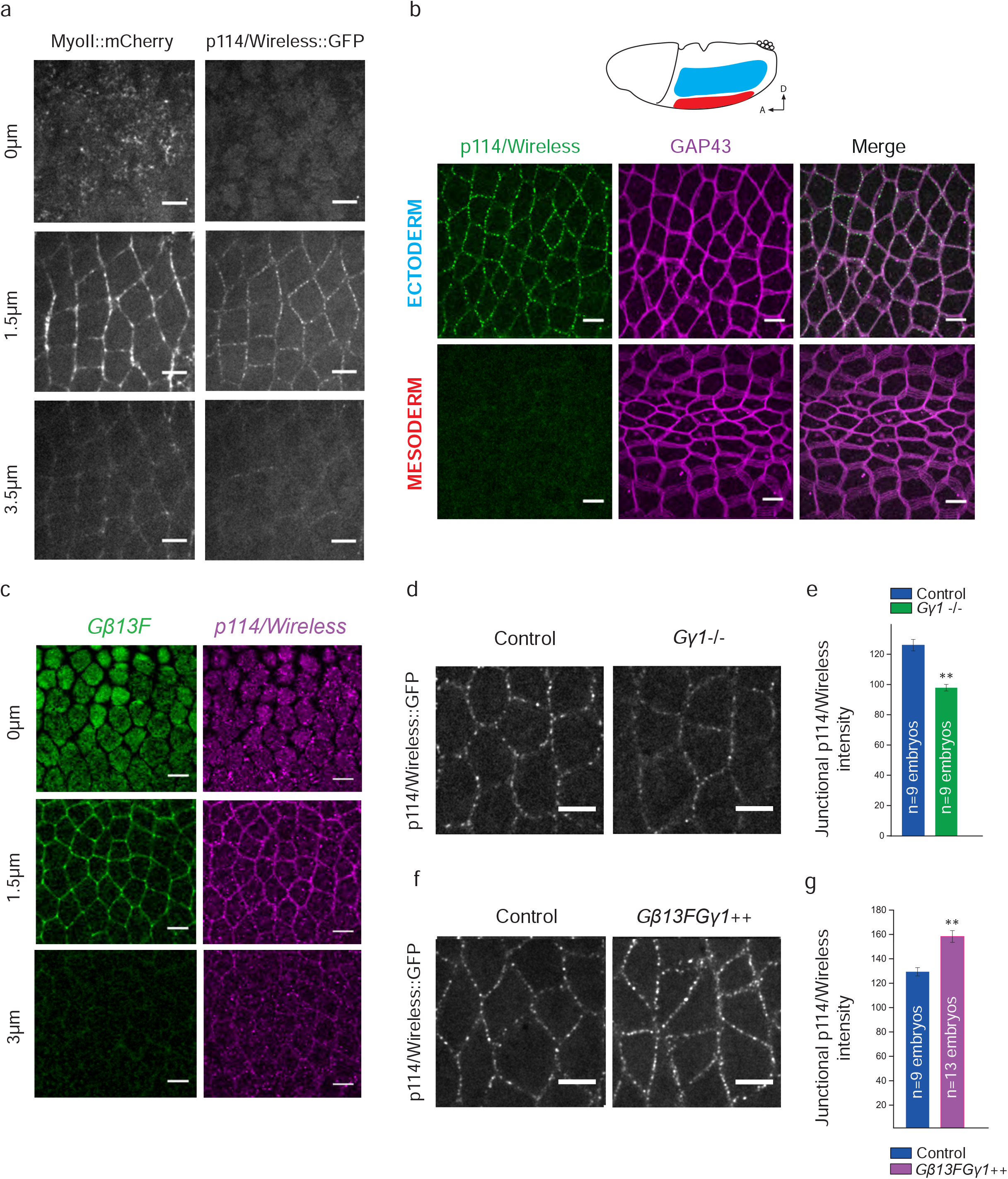
p114RhoGEF/Wireless localizes at adherens junctions under control of Gβ13FGγ1 in the ectoderm. (a) Apical (0µm), junctional (1.5µm) and lateral (3.5µm) confocal z-sections of ventro-lateral ectodermal cells from embryos co-expressing MyoII::mCherry and p114RhoGEF/Wireless::GFP. p114RhoGEF/Wireless localizes exclusively at junctions together with junctional Myo-II. (b) Confocal acquisitions of ectodermal cells (top panels) and mesodermal cells (bottom panels) in embryos expressing p114RhoGEF/Wireless::GFP and GAP43::Cherry. While p114RhoGEF/Wireless::GFP is detected at junctions in the ectoderm, p114RhoGEF/Wireless::GFP signal is absent from the invaginating mesoderm. (c) Anti-p114RhoGEF/Wireless::GFP and anti-Gβ13F stainings in ectodermal cells showing the enrichment of both p114RhoGEF/Wireless and Gβ13F at adherens junctions (1.5µm single-plane). (d, f) Confocal z-projections of ectodermal cells expressing p114RhoGEF/Wireless::GFP in control, Gγ1^-^ and Gβ13FGγ1++ embryos. Junctional p114RhoGEF/Wireless is decreased in Gγ1 germline clones and increased upon Gβ13FGγ1 overexpression. (e, g) Quantifications of mean junctional p114RhoGEF/Wireless::GFP intensities in control, Gγ1^-^ and Gβ13FGγ1++ embryos. Scale bars = 5µm. Means ± SEM are shown. Statistical significance has been calculated using Mann-Whitney U test. ns, p>0.05; * p<0.05; ** p<0.01.

Planar polarized Rho1 activity at ectodermal junctions could be explained by a planar polarized distribution of its direct activator(s) in the ectoderm. To test this hypothesis, we next compared junctional p114RhoGEF/Wrl distribution with the distribution of the non-polarized membrane protein GAP43 in the ectoderm of the same embryos. No difference was observed between p114RhoGEF/Wrl and GAP43 amplitude of polarity (Fig. S5 b and c). Thus, p114RhoGEF/Wrl localization alone cannot account for the polarized Rho signaling at junctions.

Alternatively, p114RhoGEF/Wrl activity could be polarized at junctions. Considering the newly uncovered genetic interaction between p114RhoGEF/Wrl and Gβ13F/Gγ1 in the control of junctional Rho1 signaling, we hypothesized that the heterotrimeric G proteins could be upstream activators of p114RhoGEF/Wrl. Thus, the localization of Gβ13F/Gγ1 could instruct planar polarization of p114RhoGEF/Wrl activity. We generated antibodies against two different peptides of Gβ13F (see Material and Methods) and confirmed their specificity by Western-blot and immunochemistry analyses (Fig.S6 a-c). Both antibodies revealed an apical and junctional enrichment of Gβ13F in the ectoderm (Fig.S6 c). Furthermore, Gβ13F co-localizes with both p114RhoGEF/Wrl and β-catenin at junctions (Fig.7 c and Fig.S6 d respectively) where it is not planar polarized (Fig.S6 e).

Finally, we asked whether Gβ13F/Gγ1 control Wireless enrichment at junctions. We looked at p114RhoGEF/Wrl::GFP signal in both gain (Gβ13F/Gγ1++) and loss of Gβ13F/Gγ1 (Gγ1-/-). p114RhoGEF/Wrl was decreased at junctions upon Gγ1 depletion (Fig.6 d and e). Conversely, Gβ13F/Gγ1 overexpression led to an increase in p114RhoGEF/Wrl::GFP levels at junctions though, strikingly without any gain in planar polarity (Fig.7 f and g, Fig.S6 f), which contrasts with the gain in Rho1-GTP and MyoII planar polarity in this condition. Taken together, our data show that Gβ13F/Gγ1 subunits are present at adherens junctions where they increase recruitment of p114RhoGEF/Wrl allowing Rho1 to signal efficiently in this compartment.

## Discussion

Critical aspects of cell mechanics are governed by spatial-temporal control over Rho1 activity during *Drosophila* embryo morphogenesis. This work sheds new light on the mechanisms underlying polarized Rho1 activation during intercalation in the ectoderm. We found that Rho1 activity is driven by two complementary RhoGEFs under spatial control of distinct heterotrimeric G protein subunits (Fig.S7). Notably, we uncovered a regulatory module specific for junctional Rho1 activation.

We identified p114RhoGEF/Wrl as a novel activator of junctional Rho1 in the extending ectoderm. Hence, two RhoGEFs, p114RhoGEF/Wrl and RhoGEF2, coordinate the modular Rho GTPase activation during tissue extension of the ectoderm. The division of labor in the molecular mechanisms of Rho1 activation in distinct cellular compartments lends itself to differential quantitative regulation. The activation kinetics of these different GEFs and nucleotide exchange catalytic efficiencies are likely to differentially impact Rho1 activity and therefore MyoII activation at the junctional and medial-apical compartments. For example, RhoGEF2 mammalian orthologs, LARG and PDZ-RhoGEF, show a catalytic activity that is two orders of magnitude higher as compared with the p114RhoGEF/Wrl orthologs subfamily (Jaiswal et al., 2013). This may help to establish specific contractile regimes of actomyosin in given subcellular compartments. It is therefore important to tightly control RhoGEFs localization and activity to ensure a proper quantitative activation of the downstream GTPase.

RhoGEF2 is a major regulator of medial-apical Rho1 activity during *Drosophila* gastrulation (Barrett et al., 1997; Häcker and Perrimon, 1998; Kölsch et al., 2007). Originally characterized in the invaginating mesoderm, we found that RhoGEF2 also activates Rho1 medial-apical activity in the elongating ectoderm. There, RhoGEF2 localizes both medial-apically and at junctions where it is also planar-polarized. Although RhoGEF2 and active Rho1 are both planar polarized at junctions, in *RhoGEF2* mutants, junctional Rho1-GTP is not affected and ectopic recruitment of RhoGEF2 following expression of Gα_12/13_^Q303L^ does not cause ectopic junctional Rho1-GTP accumulation. Thus, RhoGEF2 localization at the membrane is not strictly indicative of its activation status. Interestingly, Gα_12/13_/Cta is necessary for RhoGEF2 to translocate from microtubules plus ends to the plasma membrane where it signals but Gα_12/13_/Cta alone does not account for the restricted activation of Rho1 medial-apically.

We hypothesize that additional factors must regulate the spatial distribution of RhoGEF2 activity. In principle, RhoGEF2 signaling activity could either be specifically induced medial-apically independent of RhoGEF2 recruitment or RhoGEF2 could be inhibited at junctions and laterally. Sequestration of inactive RhoGEFs at cell junctions has been reported previously in mammalian cell cultures (Aijaz et al., 2005; Terry et al., 2012), suggesting that such mechanism could be evolutionary conserved. Phosphorylation can control the activity of the RH-RhoGEFs subfamily (Chikumi et al., 2002; Suzuki et al., 2003).Therefore, phosphorylation could promote activation or inhibition of RhoGEF2 activity in specific subcellular compartments in the ectoderm. RhoGEF2 is reported to be phosphorylated in the gastrulating embryo (Sopko et al., 2014).

Complementary to RhoGEF2, p114RhoGEF/Wireless is the main activator of junctional Rho1 in the ectoderm. p114RhoGEF/Wrl strictly localizes at junctions (Fig.7 a), providing a direct explanation for its junctional specific effect. We showed that Gβ13F/Gγ1 is also enriched at adherens junctions where it controls p114RhoGEF/Wrl junctional recruitment (Fig.7 d-g). Therefore, we suggest that Gβ13F/Gγ1-dependent tuning of junctional Rho1 activation could be achieved through its ability to concentrate the GEF at junctions. Gβ/Gγ–dependent regulation of RhoGEFs has been described in mammals (Niu et al., 2003a; Wang et al., 2009). One study proposes that mammalian p114RhoGEF, may bind and be activated by Gβ1/Gγ2 (Niu et al., 2003b). Interestingly, recent work demonstrates that Gα_12_ can also recruit p114RhoGEF at cell junctions under mechanical stress in mammalian cell cultures where it promotes RhoA signaling (Nestor-bergmann et al., 2018). However, the region of mammalian p114RhoGEF that binds to Gα_12_ is absent in invertebrate RhoGEFs (Martin et al., 2016). Therefore, a specific Gα_12_ control of p114RhoGEF may have appeared during vertebrate evolution while Gβ13F/Gγ1 controls p114RhoGEF ortholog in invertebrates. How Gβ13F/Gγ1 control p114RhoGEF/Wrl at junctions in the *Drosophila* embryo remains an open question.

Importantly, neither Gβ13F/Gγ1 nor p114RhoGEF/Wrl are themselves planar polarized at junctions. Hence, their distribution alone cannot explain polarized Rho1 activity at junctions. However, our Gβ13F antibodies detect both inactive Gβ13F (bound to Gα-GDP) and active Gβ13F (released from Gα-GTP) and it would be therefore important to probe specifically the distribution of active Gβ13F/Gγ1 dimers in the ectoderm. Strikingly we found that an increase in Gβ13F/Gγ1 dimers hyperpolarizes Rho1 activity and Myo-II at vertical junctions (Fig.5h). Gβ13F/Gγ1 overexpression also leads to an overall increase in p114RhoGEF/Wrl levels at junctions, although p114RhoGEF/Wrl is not planar polarized in this condition. This indicates that recruitment at the plasma membrane and activation of p114RhoGEF/Wrl are independently regulated, similar to RhoGEF2. In contrast, p114RhoGEF/Wrl overexpression increases Myo-II at both transverse and vertical junctions, although a slightly stronger accumulation is observed at vertical junctions (Fig.4 j, k). Therefore, while p114RhoGEF/Wrl junctional levels are increased in both experiments, only Gβ13F/Gγ1 overexpression leads to an increased planar polarization of Rho1-GTP and Myo-II at vertical junctions. This points to a key role for Gβ13F/Gγ1 subunits in the planar-polarization process associated with, but independent from the sole recruitment of p114RhoGEF/Wrl at junctions. In principle, Gβ13F/Gγ1 could bias junctional Rho1 signaling either by promoting its activation at vertical junctions or by inhibiting it at transverse junctions. Gβ13F/Gγ1 could also control active Rho1 distribution independent of its activation. For instance, a scaffolding protein binding to Rho1-GTP at junctions could be polarized by Gβ13F/Gγ1 to bias Rho1-GTP distribution downstream of its activation. Anillin, a Rho1-GTP anchor known to stabilize Rho1 signaling at cell junctions (Budnar et al., 2018), is a potential candidate in the ectoderm. Last, Toll receptors control Myo-II planar-polarity in the ectoderm (Paré et al., 2014). Whether Gβ13F/Gγ1 and Tolls are part of the same signaling pathway is an important point yet to address in the future.

Finally, our study sheds light on new regulatory differences underlying tissue invagination and tissue extension. Here we found that p114RhoGEF/Wrl localizes at junctions in the ectoderm where it activates Rho1 and Myo-II. In contrast, maternally and zygotically supplied p114RhoGEF/Wrl::GFP is not detected at junctions in the mesoderm. We see little if any cytoplasmic signal in this condition suggesting that p114RhoGEF/Wrl::GFP could be degraded in these cells. Interestingly, the E3 ubiquitin-ligase Neuralized is active only in the mesoderm where it controls the invagination of the tissue (Perez-Mockus et al., 2017). Whether p114RhoGEF/Wrl is a target of Neuralized in the mesoderm is unknown. Since p114RhoGEF is expressed maternally this mechanism would ensure absence of this protein in the mesoderm despite maternally loaded mRNA. Additionally, virtual *in situ* hybridizations suggest that p114RhoGEF/Wrl mRNA is not zygotically expressed in the mesoderm but is present in the rest of the embryo (Karaiskos et al., 2017). Thus, repression of both p114RhoGEF/Wrl mRNA and protein in the mesoderm could be an important mechanism for cell apical constriction and proper tissue invagination. Of interest, Rho1 signaling is absent at junctions in the mesoderm (Mason et al., 2013). Therefore, it is tempting to suggest that the absence of p114RhoGEF/Wrl at junction in the mesoderm accounts for cells inability to activate Rho1 in this compartment. Importantly, the GPCR Smog and Gβ13F/Gγ1 subunits, found to control junctional Rho1 in the ectoderm, are present to both tissues (Kerridge et al., 2016). p114RhoGEF/Wrl differential expression and/or subcellular localization could be a key element to bias signaling towards junctional compartment in the ectoderm.

Cell contractility necessitates activation of the Rho1-Rock-MyoII core pathway. During epithelial morphogenesis, tissue and cell-specific regulation of Rho1 signaling requires the diversification of Rho1 regulators, in particular RhoGEFs and RhoGAPs, as shown in this study. There are at least 26 RhoGEFs and 23 RhoGAPs encoded in the *Drosophila* genome. Some of them are tissue specific with given subcellular localizations and activation mechanisms. The identification of signaling modules, namely Gα_12/13_-RhoGE F2 and Gβ13F/G γ1-p114RhoGEF/Wrl, provides a simple mechanistic framework for explaining how tissue specific modulators control Rho1 activity in a given subcellular compartment in a given cell type. Therefore, we suggest that the variation of (1) ligands, GPCRs and associated heterotrimeric G proteins, and (2) types of RhoGEFs and RhoGAPs as well as their combination, activation and localization by respective co-factors underlie the context-specific control of Rho1 signaling during tissue morphogenesis. How developmental patterning signals ultimately control Rho regulators is an exciting area for future investigations.

## Methods

### Fly stocks and genetics

The following mutant chromosomes were used: FRTG13 Gγ1^N159^ (refs Izumi et al., 2004; Kanesaki et al., 2013), FRTG13 RhoGEF2^l(2)04291^(ref Häcker and Perrimon, 1998), UAS-TRIP RhoGEF2 (Bloomington 34643), UAS-TRIP CG10188 (Bloomington 41579), UAS-TRIP Gα_12/13_ (Bloomington 51848), UAS-TRIP Yellow (Bloomington 64527), pUASt-Gα_12/13_^Q303L^ (ref(Fuse et al., 2013), UASt-Gγ1^#15^ (ref Kanesaki et al., 2013), UASt-Gβ13F^#20^ (ref Kanesaki et al., 2013), Rho1-mCh::Rho1 (ref Abreu-Blanco et al., 2014), Ubi-Ani-RBD::GFP (ref Munjal et al., 2015), UAS-EB1::GFP (Gift from Brunner, ref Jankovics and Brunner, 2006), UASp-RFP::RhoGEF2 (ref Wenzl et al., 2010), RhoGEF2-GFP::RhoGEF2 (ref Mason et al., 2016), UASp-mCh::GAP43 (gift from Manos Mavrakis) and endo-α-Catenin::YFP (Cambridge Protein Trap Insertion line (CPTI-002516); DGRC #115551). endoCAD::GFP replaces endogenous E-cadherin protein at the locus (ref Huang et al., 2009) and sqh_RLC-Myosin-II::mCherry (chromosome 2 or 3; ref Martin et al., 2009 + chromosome 2; ref Bailles et al., 2018). 67-Gal4 (mat-4-GAL-VP16), nos-Gal4 and 15-Gal4 are ubiquitous, maternally supplied, Gal4 drivers. Germline clones for Gγ1^N159^ and RhoGEF2^l(2)04291^ were made using the FLP-DFS system (refs Chou and Perrimon, 1996). All fly constructs and genetics are listed in Supplementary Extended Table 1 and Table 2.

### Transgenic lines

SqhPa-p114/Wireless expression vectors were generated using a SqhPa-sqh::mCherry modified vector (kind gift from A. Martin), a pCasper vector containing a sqh (MyoII RLC, CG3595) minimal promoter. A PhiC31 attB sequence was inserted downstream from the white gene of the SqhP vector into AfeI restriction site to perform PhiC31 site specific transgenesis. To build SqhPa-p114/Wireless plasmids, ORF of sqh::mCherry was replaced by the one of p114/Wireless (CG10188) using 2 ESTs as matrices (RE42026 and RE33026) to build a WT sequence (Genebank, NP_609977). p114/Wireless was then tagged either N- or C-terminally by mEGFP with a SGGGGS flexible aa linker in between. SqhPa-p114/Wireless (CG10188) - shRNA^R^ Resistant was built by introducing silent point mutations to the codons of the 21bp targeted by the shRNA TRIP 41579 (CACGAGACAGACAATGGATTA to CAtGAaACtGAtAAcGGtTTA). All recombinant expression vectors were verified by sequence (Genewiz) and were sent to BestGene Incorporate for PhiC31 site specific mediated insertion into attP2 (3L, 68A4). FASTA sequences of these vectors are available on request.

### Antibody generation

To generate specific antibodies for Gβ13F, peptides corresponding to the amino-terminal region and internal region of the Gβ13F protein were commercially synthesized and used to immunize rabbits (Eurogentec). The peptide sequences employed were as follows: MNELDSLRQEAESLK (aa 1-15) and CKQTFPGHESDINAVT (aa 218-233). Polyclonal anti-Gβ13F antibodies affinity purified against the immunizing peptide were then tested for specificity in western blots and immunostainings. Lysates from dechorionated embryos were prepared in 10 mM Tris/Cl pH 7.5; 150 mM NaCl; 0.5 mM EDTA; 0.5% NP-40 supplemented with HALT Protease/Phosphatase Inhibitor Mix (Life Technologies) and 0.2M PMSF (Sigma). Samples were denatured, reduced, separated by SDS PAGE and transferred to PVDF membranes. After blocking, blots were incubated with polyclonal antibody (2µg/mL) with or without preincubation of antibody with 200 µg/ml of immunizing/affinity purified peptide. A band of the expected molecular weight (43 kD) was present in the western blot and was abolished when the antibody was preincubated with the immunizing peptide. Similarly, the signal observed in subsequent immunofluorescence labelings was abolished when the antibody was preincubated with the immunizing peptide.

### Immunofluorescence

Methanol-heat fixation(H.-Arno J. Müller and Eric Wieschaus, 1996) was used for embryos labeled with rabbit anti-Gβ13F (1:20, as described above), mouse anti-β catenin (1:100, DSHB), mouse anti-Neurotactin (1:50, DSHB). A chicken anti-GFP antibody (1:1000, Aves Labs) was used in embryos expressing GFP::Wireless to amplify the signal in fixed embryos. Secondary antibodies (Invitrogen) were used at 1:500. Fixed samples (using Aqua-Poly mount, Polysciences) were imaged under a confocal microscope (LSM 780, Carl Zeiss) using a Plan Apochromat 40x/1.4 NA oil immersion objective.

### Differential interference contrast live imaging

Standard techniques were used to immobilize embryos for imaging. Bright-field time-lapse images were collected on an inverted microscope (Zeiss) and a programmable motorized stage to record different positions over time (Mark&Find module from Zeiss). The system was run with AxioVision software (Zeiss) and allowed the acquisition of time-lapse data sets in wild-type or mutant embryos. Images were acquired every 2 min for 40 min post dorsal movement of the posterior pole cells. The extent of elongation was measured by tracking the distance between the pole cells and the posterior pole at each time point and normalized to the total length of the embryo.

### Embryo viability test

40 freshly hatched females and males were incubated at 25°C for 4 days in each experimental conditions (Control, p114/Wireless shRNA and p114/Wireless shRNA, Sqh-p114/Wireless shRNA^R^). For egg collection, flies were given a fresh apple juice agar plate to lay eggs for 4 hours. Eggs were then counted and incubated at 25°C for 2 days. The total number of emerging larva was counted and plotted in percentage as a function of viability.

### Embryo injection

Microtubule depolymerization was carried out by injecting Colcemid (500µg/ml in water, 234109-M, Sigma-Aldrich) in y[*] w[67c23] or EndoCad::GFP, Sqh::mCh embryos during the fast phase of cellularization. Subsequently, embryos were filmed at the onset of germ-band extension on a Nikon Roper spinning disc Eclipse Ti inverted microscope using a 100X_1.4 N.A. oil-immersion objective or on a Zeiss inverted bright field microscope.

### Image acquisition

Embryos were prepared as described before (Pilot et al, 2006). Timelapse imaging was done from stage 6 during 15 to 30 min depending on the experiment, on a Nikon Roper spinning disc Eclipse Ti inverted microscope using a 100X_1.4 N.A. oil-immersion objective or a 40X_1.25 N.A. water-immersion (for cell-intercalation measurement) at 22°C. The system acquires images using the Meta-Morph software. For medial and junctional intensity measurements, 10 to 18 Z sections (depending on the experimental conditions), 0.5µm each, were acquired every 15s. Laser power was measured and kept constant across all experiments.

### Image analysis

All image processing was done in imageJ/Fiji free software. For all quantifications for medial and junctional Rho1-GTP and Myo-II, maximum-intensity z-projection of slices was used, followed by a first background subtraction using the rolling ball tool (radius 50 pixels∼4µm) and a second subtraction where mean cytoplasmic intensity value measured on the projected stack was subtracted to the same image. Cell outlines were extracted from spinning disk confocal images of Ecad::GFP or Rho1-GTP using the Tissue Analyzer software (Aigouy et al., 2010) from B.Aigouy (IBDM, France). The Ecad-GFP resulting outlines were then dilated by 2 pixels on either side of the junction (5-pixel-wide lines) and used as a junctional mask on the MyoII::mCherry channel. Medial-apical area was obtained by shrinking individual cell mask by 4 pixels to exclude any contribution of junctional signal (ImageJ/Fiji macro, Girish Kale, IBDM France). Medial and junctional Myo-II and Rho1-GTP values were mean intensities calculated in these two non-overlapping cell areas.

For planar polarity analysis, junctional masks described previously were used to extract for each junction the mean pixel intensity and orientation. Intensities were averaged for all junctions in each angular range. Amplitude of polarity was then calculated as a ratio between signal intensity measured at vertical junctions (75-90° angular range) over intensity measured at transverse junctions (0-15° angular range).

To measure the number of T1 transitions, Tissue Analyzer software. Segmentation was automatically performed on Utr::GFP channel by the plugin and corrected by the experimenter. Tracked cells present in the field of view during a period of 10min were then analyzed for T1 events. T1 events were automatically detected by the plugin and checked manually to prevent false detections.

### Statistics

Errors bars are SEM unless otherwise indicated. Statistical significance was determined and *P* values calculated with a non-parametric Mann–Whitney *U* test or a Kolmogorov-Smirnov two-sample test in Origin (v8). The experiments were not randomized, and the investigators were not blinded to allocation during experiments and outcome assessment.

## Acknowledgement

We are grateful to N.Fuse (Kyoto, Japan), F.Matsuzaki (RIKEN, Japan), A.Martin (MIT, USA), J.Großhans (Institut of Developmental Biochemistry Göttingen, Germany), S.Kerridge (IBDM, France), the Drosophila Genetic Resource Center, and the Bloomington Stock Center for the gift of flies. We thank the TRiP at Harvard Medical School (NIH/NIGMS R01-GM084947) for providing transgenic RNAi fly stocks used in this study. We thank members of the Lecuit group and C.P.Toret (IBDM,France) for stimulating discussions and comments during the course of this project and writing of this manuscript. We thank B.Dehapiot (IBDM,France), G.Kale and C.Collinet (IBDM,France) for help with cell segmentation/tracking and image processing. This work was supported by the ERC (Biomecamorph no. 323027) and the Ligue Contre le Cancer. A.G.D.L.B was supported by the Ministère de l’Enseignement supérieur, de la Recherche et de l’Innovation and the Fondation Bettencourt Schueller. We also acknowledge the France-BioImaging infrastructure supported by the Agence Nationale de la Recherche (ANR-10-INSB-04-01, call ‘Investissements d‘Avenir’).

## Author contributions

A.G.D.L.B and T.L conceived the project. A.G.D.L.B and T.L analyzed the data. A.G.D.L.B performed all the experiments and data analysis except for Supplementary Fig.S7 b performed by A.C.L. J-M.P created all the constructs and performed all cloning and molecular characterization. A.G.D.L.B and T.L wrote the paper. All authors commented on the manuscript.

---

**Figure S1.**
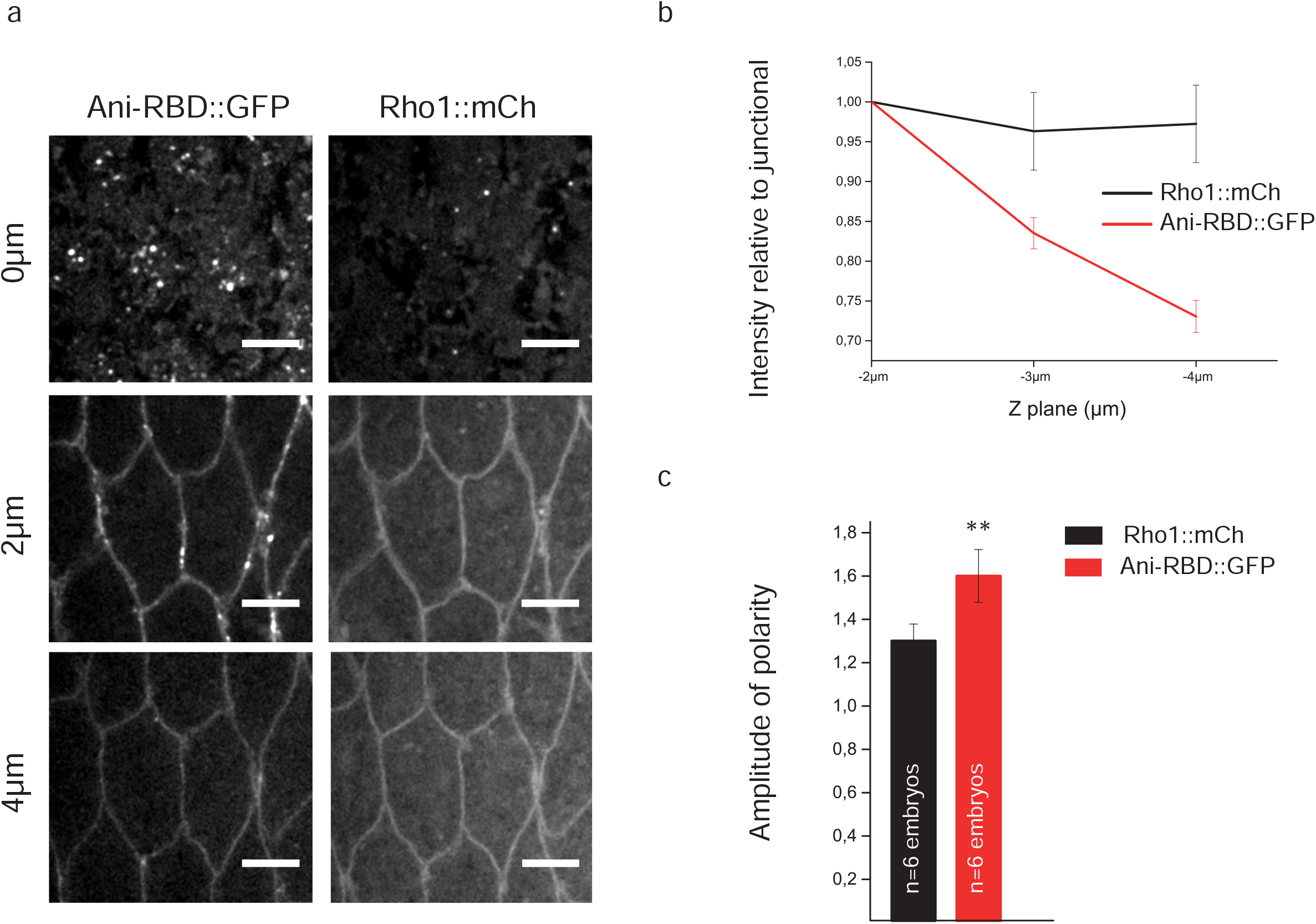
Rho1 protein is uniformly distributed in the ectoderm while its activity is polarized. (a) Apical (0µm), junctional (2µm) and lateral (4µm) confocal z-sections of ectodermal cells co-expressing Ani-RBD::GFP and Rho1::mCherry. Active Rho1 is enriched medial-apically and at junctions where it is planar-polarized. Rho1::mCh signal is homogenous along the apico-basal axis. (b) Ani-RBD::GFP and Rho1::mCherry cortical levels normalized to the apical junctional intensities (−2µm below the apical membrane) along the apico-basal axis. While Rho1::mCh signal is uniform at cell cortex along the antero-posterior axis, active Rho1 is specifically enriched apically. (c) Quantification of Rho1::mCherry and Ani-RBD::GFP amplitude of polarity at junctions in the same embryos. A higher amplitude of polarity is measured for Ani-RBD::GFP at junctions compared to total Rho1. All the panels have the same orientation: dorsal at the top, anterior to the left. Scale bars = 5µm. Means ± SEM are shown. Statistical significance has been calculated using Mann-Whitney U test. ns, p>0.05; * p<0.05; ** p<0.01.

**Figure S2.**
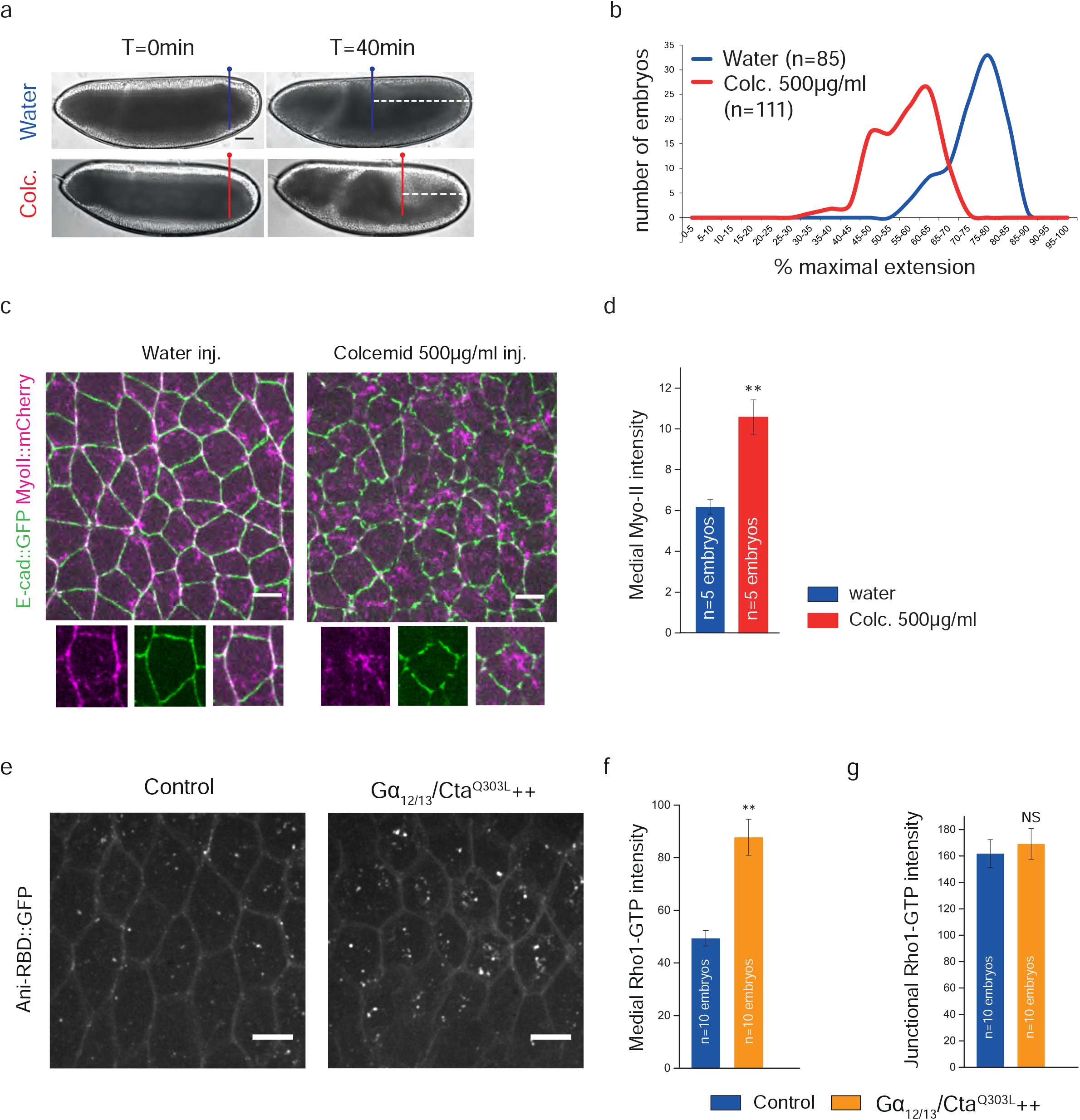
Microtubule depolymerization and Gα_12/13_/Cta^Q303L^ overexpression increases medial-apical Rho1 signalling. (a) Lateral view of a water- and a colcemid-injected embryo at the onset (t=0min) of germ-band extension and 40min later. The dotted lines mark the distance between the pole cells and the posterior side of the embryos 40 minutes after the onset of germ-band extension. (b) Quantification of germ-band extension 40min after the onset of the process in water and colcemid-injected embryos. n=number of embryos. (c) Confocal acquisitions of water- and colcemid-injected embryos co-expressing Myo-II::mCherry and Endocad (E-cad)::GFP. A closeup of a representative cell is shown in the bottom part for both conditions. Colcemid-treated cells display higher medial-apical Myo-II levels and increased contractility. (d) Quantifications of mean medial-apical Myo-II intensities in both water- and colcemid-injected embryos. (e) 4µm confocal z-projection of ventro-lateral ectodermal cells expressing Ani-RBD::GFP in control and Gα_12/_13/CtaQ303L++ embryos. Active Rho1 is specifically increased in the medial-apical compartment of the cells upon Gα_12/_13/CtaQ303L overexpression. (f, g) Mean medial-apical and junctional Rho1-GTP intensities in control and Gα_12/_13/CtaQ303L++ embryos. All the panels have the same orientation: dorsal at the top, anterior to the left. Scale bars = 50µm (a) and =5µm (c and e). Means ± SEM are shown. Statistical significance has been calculated using Mann-Whitney U test. ns, p>0.05; * p<0.05; ** p<0.01.

**Figure S3.**
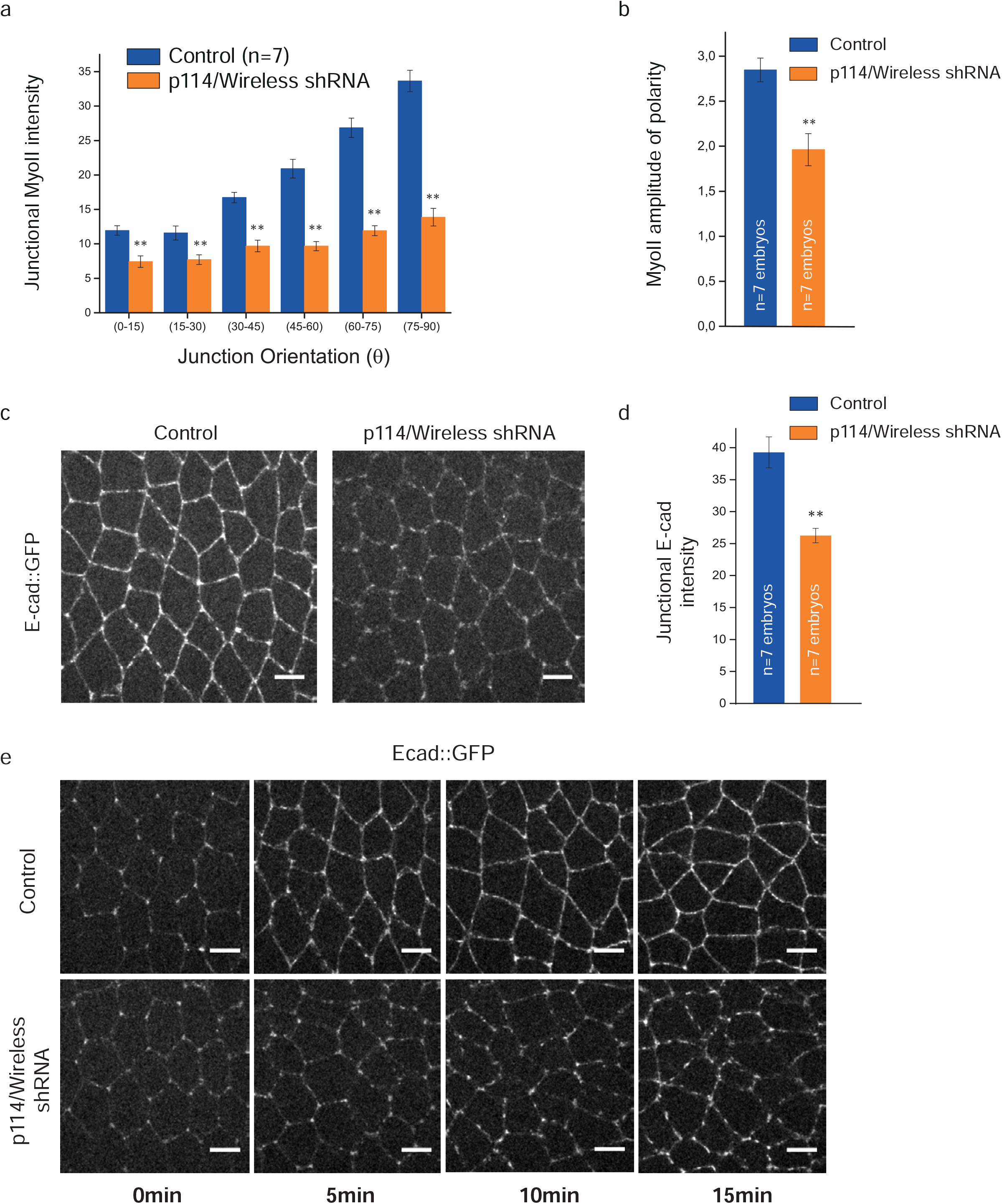
Myo-II and E-cadherin junctional levels are affected in p114RhoGEF/Wireless knock-down embryos. (a) Mean junctional intensity of Myo-II according to the angle of the junctions in control and p114Rhogef/Wireless shRNA expressing embryos. (junction angle; 0°, parallel to the antero-posterior axis; 90°, perpendicular to the antero-posterior axis). n= number of embryos. A global decrease in Myo-II is observed at both transverse and vertical interfaces. (b) Quantification of junctional Myo-II amplitude of polarity in control and p114Rhogef/Wireless shRNA embryos. A reduction of Myo-II polarity is observed upon p114Rhogef/Wireless knock-down. (c) Confocal projections of ectoderm tissues expressing E-cad::GFP in control and p114Rhogef/Wireless shRNA embryos. E-cadherin junctional levels are decreased upon p114/Wireless depletion. (d) Mean E-cadherin junctional intensities. (e) E-cad::GFP in time lapse videos of control (top panels) and p114Rhogef/Wireless shRNA embryos (bottom panels) (t=0 is the end of the mesoderm pulling). Anterior is left and ventral is down. E-cadherin is enriched at cell vertices in the early germ-band and rapidly accumulates along junctions where it forms an adhesive belt in control embryos. In p114Rhogef/Wireless shRNA embryos, E-cadherin junctional maturation is disrupted and shows a low and very discontinuous signal. Scale bars = 5µm. Means ± SEM are shown. Statistical significance has been calculated using Mann-Whitney U test. ns, p>0.05; * p<0.05; ** p<0.01.

**Figure S4.**
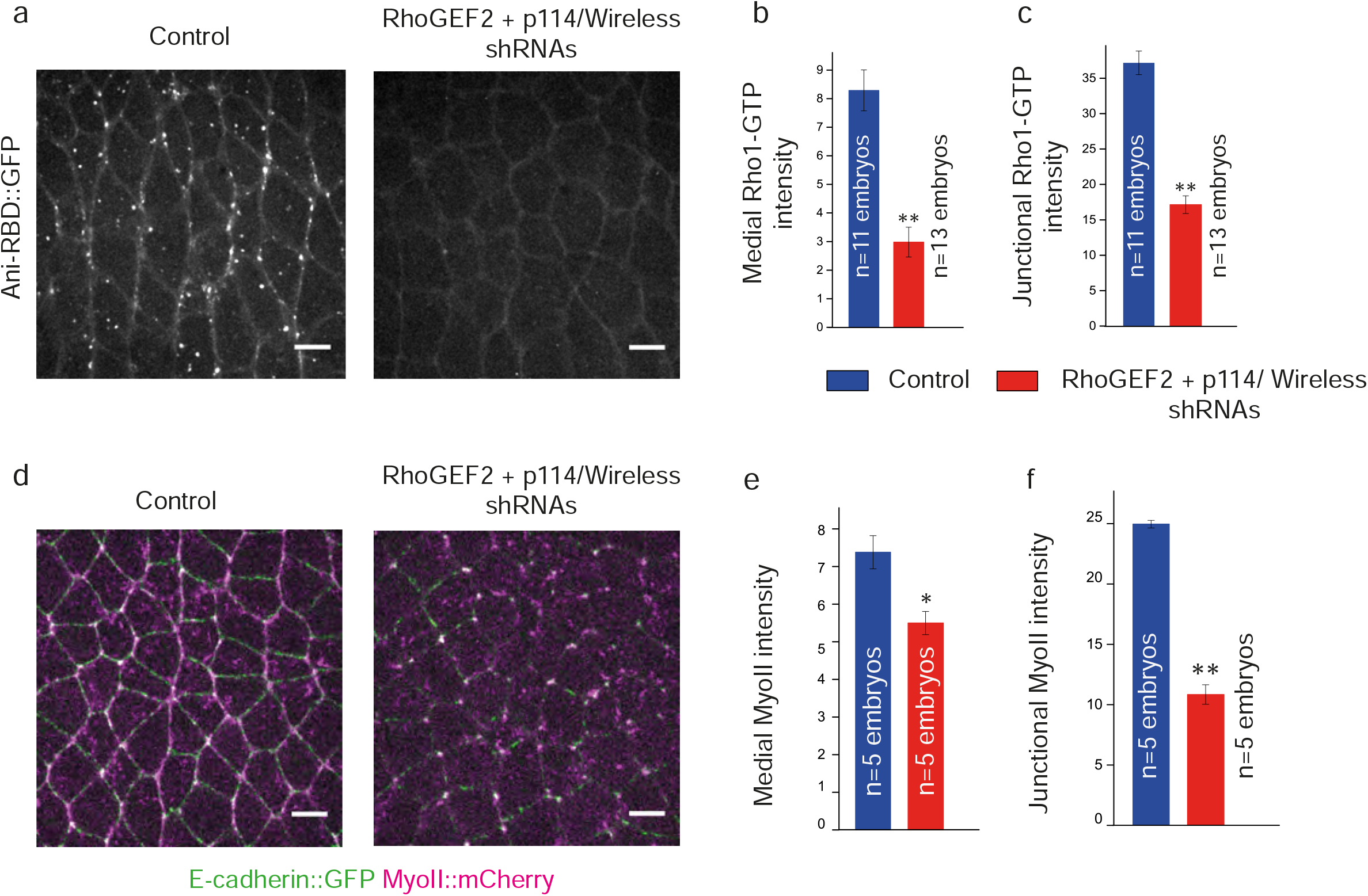
RhoGEF2 and p114RhoGEF/Wireless double knock-down decreases both medial-apical and junctional Rho1 signaling. (a) Confocal z-projections of control and RhoGEF2 + p114RhoGEF/Wireless double knock-down embryos (RhoGEF2+Wireless shRNAs) expressing Ani-RBD::GFP. A decrease in both medial-apical and junctional Rho1 activity is observed in the second condition. (b, c) Mean medial-apical and junctional Rho1-GTP intensities in control and RhoGEF2+Wireless shRNAs embryos. (d) 5µm confocal z-projections of ventro-lateral ectodermal cells expressing E-cad::GFP and MyoII::mCherry in control and RhoGEF2+Wireless shRNAs embryos. Similar to the previous observations, medial-apical and junctional Myo-II pools are decreased in mutant embryos. (e, f) Mean medial-apical and junctional Myo-II intensities. Scale bars = 5µm. Means ± SEM are shown. Statistical significance has been calculated using Mann-Whitney U test. ns, p>0.05; * p<0.05; ** p<0.01.

**Figure S5.**
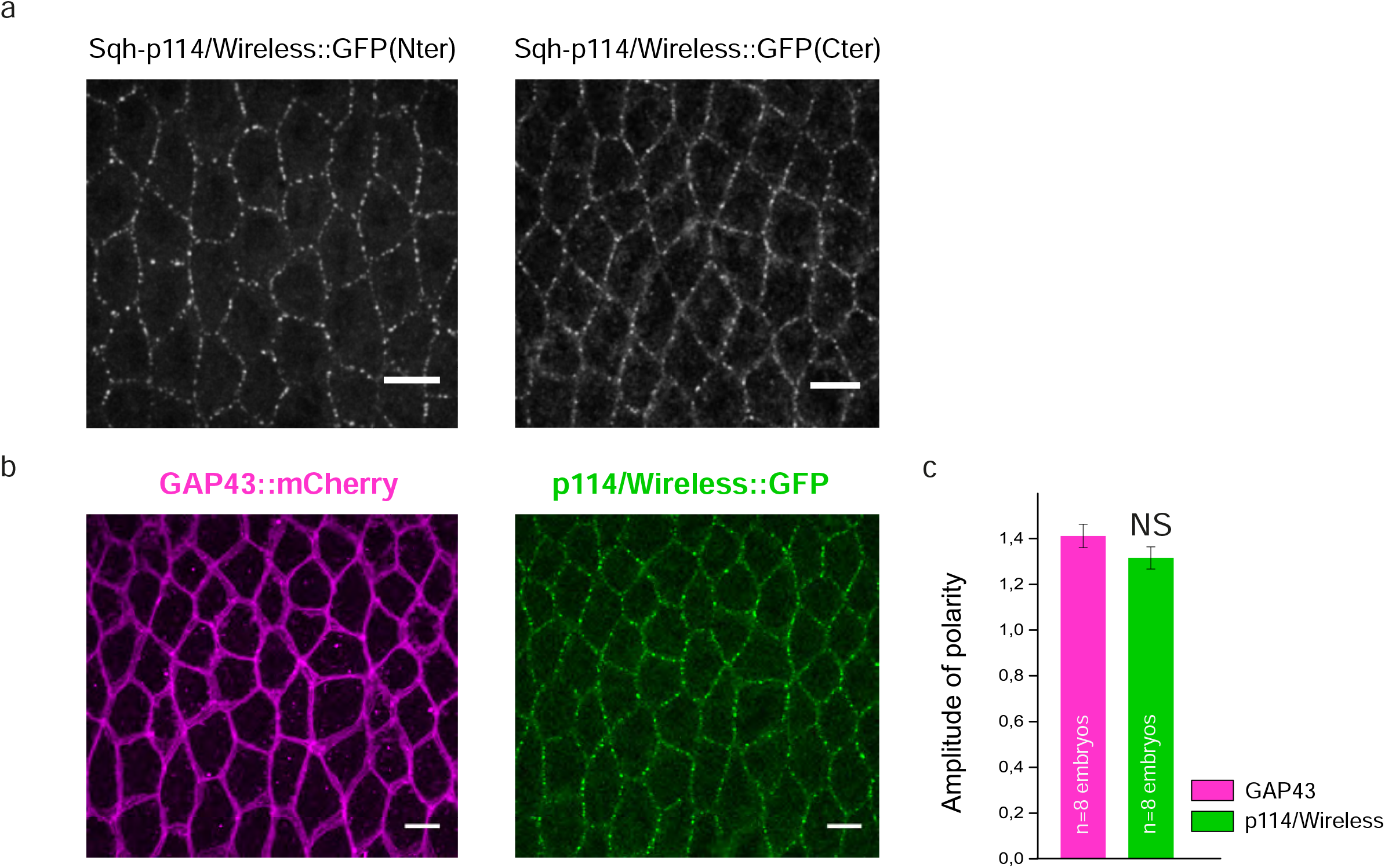
GFP-tagged p114RhoGEF/Wireless localizes at cell junctions with no apparent planar-polarity. (a) Confocal z-projections of ectodermal cells expressing p114RhoGEF/Wireless tagged with GFP at its N-terminal (left panel) or C-terminal end (right panel). Although both fusion proteins localize at adherens junctions, a stronger cytoplasmic signal is often observed in embryos expressing the p114RhoGEF/Wireless construct tagged in C-ter. p114RhoGEF/Wireless tagged in N-ter has been used hereafter. (b) 4µm confocal z-projection of ectodermal cells co-expressing the membrane marker GAP43::mCherry and p114RhoGEF/Wireless::GFP in the same embryo. (c) Quantification of GAP43 and p114RhoGEF/Wireless amplitude of polarity in the same embryos. p114RhoGEF/Wireless::GFP polarity at junctions is similar to the polarity of the membrane marker. Scale bars = 5µm. Means ± SEM are shown. Statistical significance has been calculated using Mann-Whitney U test. ns, p>0.05; * p<0.05; ** p<0.01.

**Figure S6.**
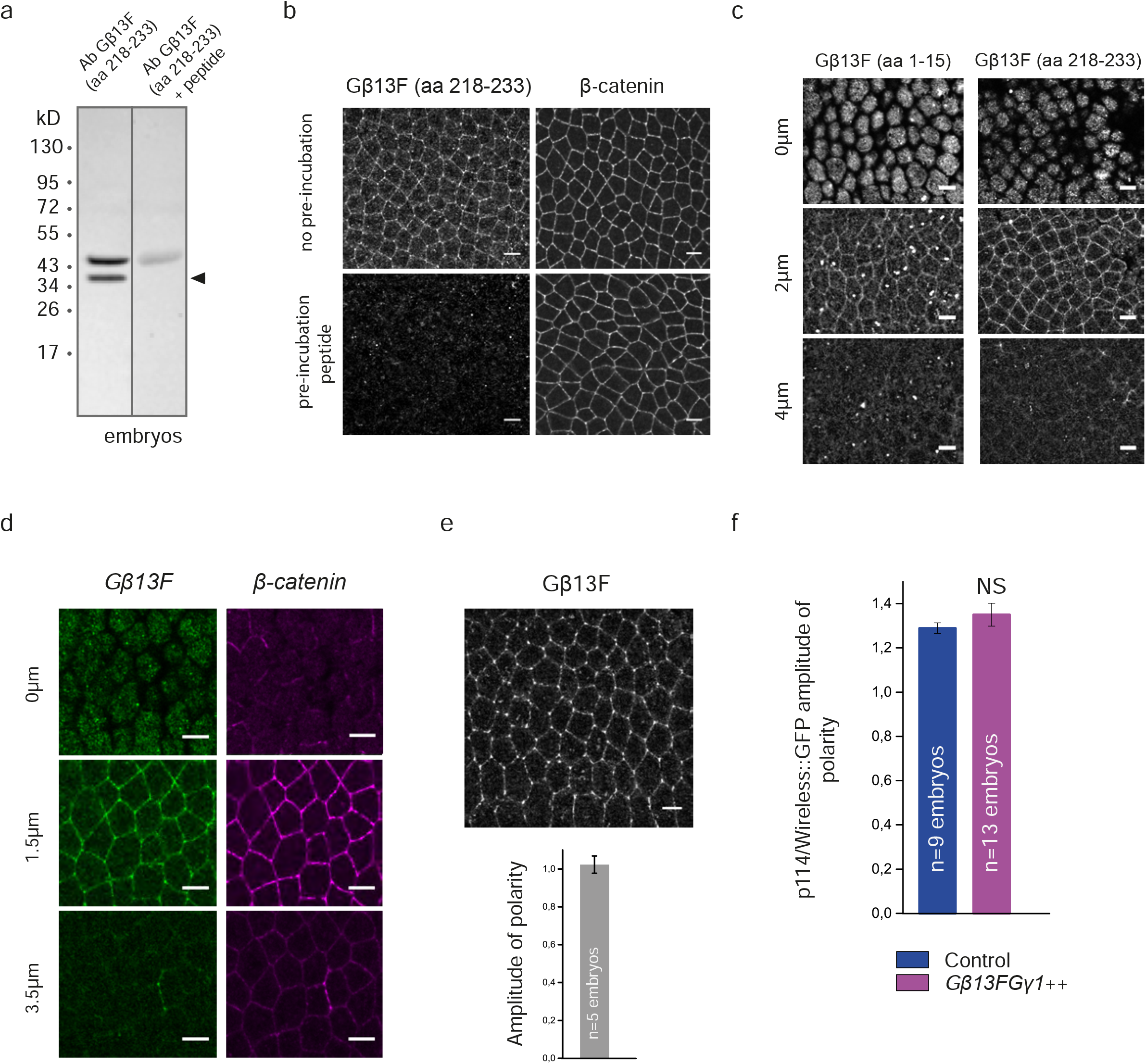
Gβ13F localizes apically and at adherens junctions with no planar-polarity in the ectoderm. (a) Gβ13F (218-233) antibody specificity of binding was analyzed further in immunoblotting on *yw* embryos lysates. Two bands were detected: one at the expected Gβ13F molecular weight (37kDa, black arrow) and another band around 45 kDa. Importantly, the 37kDa band was abolished when the membrane was pre-treated with the blocking Gβ13F (218-233) peptide. The higher molecular weight band was strongly diminished but not completely removed. (b) Ventro-lateral ectoderms stained with Gβ13F (218-233) and β-catenin antibodies. Pre-incubation of the Gβ13F (218-233) antibody with the Gβ13F (218-233) peptide completely abolished Gβ13F signal (left bottom panel). (c) Apical (0µm), junctional (2µm) and lateral (4µm) confocal z-sections of ectodermal cells in fixed embryos stained with two different purified antibodies against Gβ13F (see material and methods). Both antibodies showed a similar staining, with Gβ13F being enriched apically and at adherens junctions. Because antibody against the Gβ13F (218-233) peptide gave a cleaner staining with less intracellular aggregates, we performed the next experiments using this purified antibody exclusively. (d) Apical (0µm), junctional (1.5µm) and lateral (3.5µm) confocal z-sections of ectodermal cells in fixed embryos stained with Gβ13F and β-catenin antibodies. Gβ13F co-localizes with β-catenin at junctions. (e) Quantification of the amplitude of polarity of Gβ13F measured on fixed embryos. Gβ13F is not planar-polarized at cell junctions. (f) Quantifications of p114RhoGEF/Wireless::GFP amplitude of polarity in control and Gβ13FGγ1 overexpressing embryos (Gβ13FGγ1++). While Gβ13FGγ1 overexpression increases p114RhoGEF/Wireless::GFP levels at cell junctions (see main Fig.7 f and g), its amplitude of polarity is not affected. Scale bars = 5µm.

**Figure S7.**
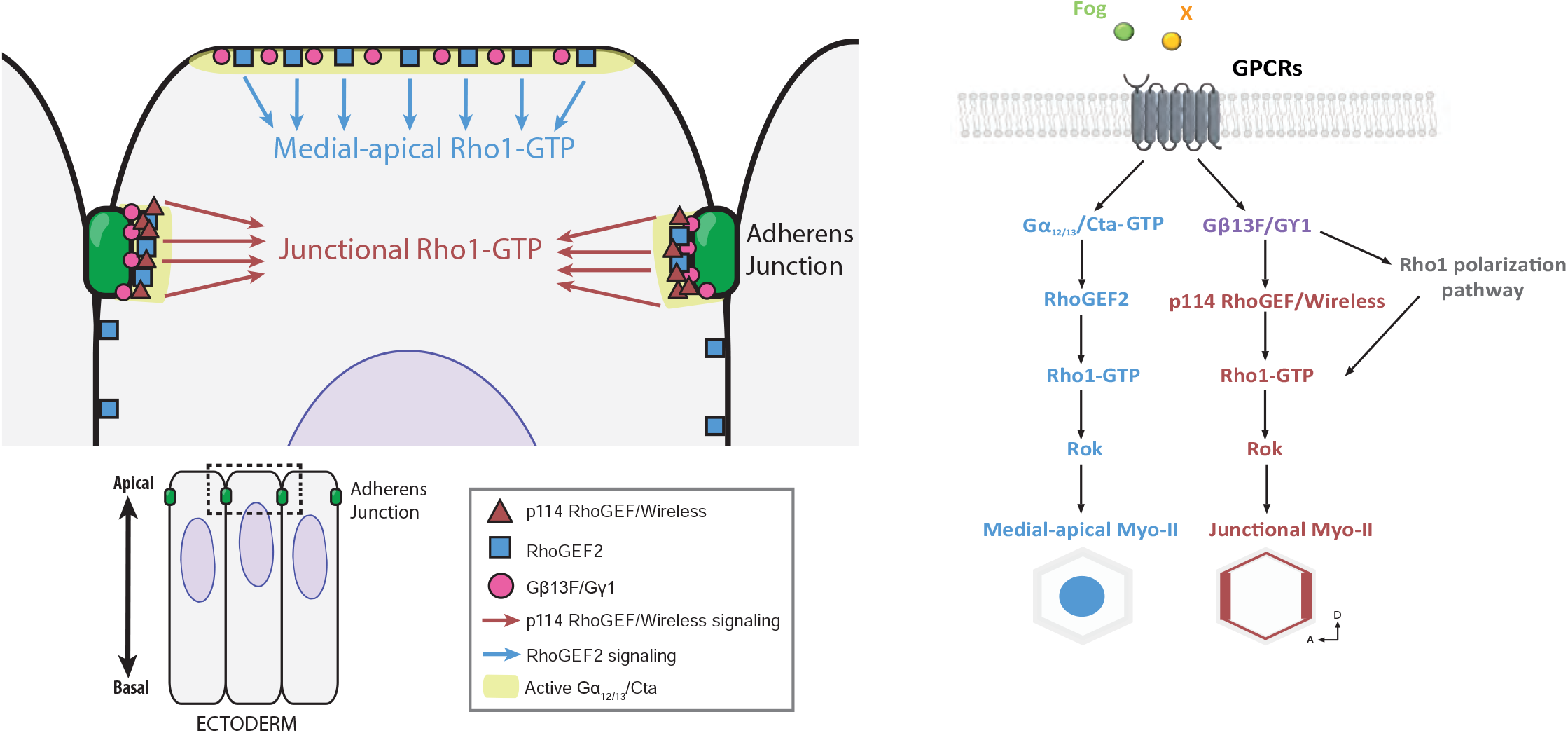
Distinct RhoGEFs compartimentalize Rho1 signaling apically and at junctions under control of G proteins in the *Drosophila* embryonic ectoderm. Left panel: a schematic view of heterotrimeric G protein subunits and RhoGEFs spatial distribution and activity in ectodermal cells. RhoGEF2 (in blue) is present apically and at adherens junctions but only activates medial-apical Rho1 signaling. Active Gα_12/13_/Cta recruits RhoGEF2 in both compartments by promoting its dissociation from microtubules. Additional regulators at the membrane bias RhoGEF2 activity towards the medial-apical compartment. p114RhoGEF/Wireless (in red) is present exclusively at junctions where it activates Rho1 under control of Gβ13FGγ1 and other unknown regulators. Right panel: An overview of the medial and junctional signaling pathways controlling Rho1 activity in the ectoderm. Following stimulation by ligand (Fog and others), GPCRs release active Gα_12/13_/Cta (Gα_12/13_/Cta-GTP) and active Gβ13FGγ1 dimers that promote RhoGEF2 and p114 RhoGEF/Wireless signaling respectively. How Gβ13FGγ1 polarize junctional Rho1 activation is unclear and could involve Toll receptors.

## Movie Legends

**Movie 1** Developing ectoderm in control and RhoGEF2 shRNA embryos. Ani-RBD::GFP, Related to Fig.1b; Scale bar=5µm. Duration: 2min

**Movie 2** EB1::GFP and RhoGEF2::RFP dynamics in an ectodermal cell from a developing embryo. Related to Fig.2b; Scale bar=5µm. Duration: 4s

**Movie 3** Developing ectoderm in control and Gα_12/13_/Cta^Q303L^++ embryos. GFP::RhoGEF2, Related to Fig.2g; Scale bar=5µm. Duration: 14s

**Movie 4** T1s events during germ-band extension in control and *CG10188* shRNA embryos. Related to Fig.3c; Scale bar=15µm. Red/Green: T1s. Duration: 10min

**Movie 5** Developing ectoderm in control and p114RhoGEF/Wireless shRNA embryos. MyoII::mCherry, Related to Fig.4d; Scale bar=5µm. Duration: 15min

**Movie 6** Developing ectoderm in control and p114RhoGEF/Wireless overexpressing embryos. MyoII::mCherry, Related to Fig.4g; Scale bar=5µm. Duration: 15min

**Movie 7** Developing ectoderm in control and Gβ13F/Gγ1 overexpressing embryos. MyoII::mCherry, Related to Fig.5g; Scale bar=5µm. Duration: 17min

---

**Supplementary table 1.**
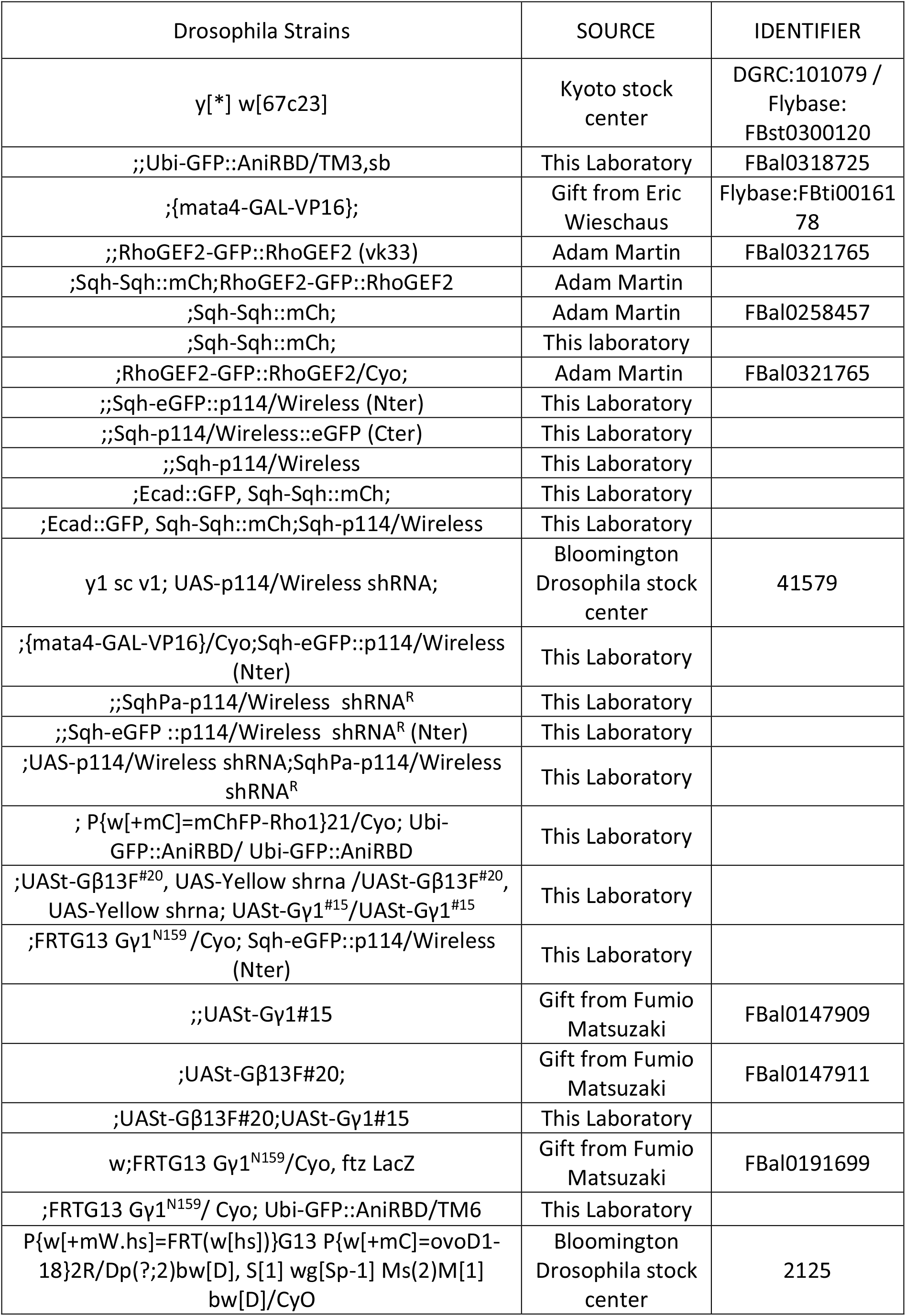
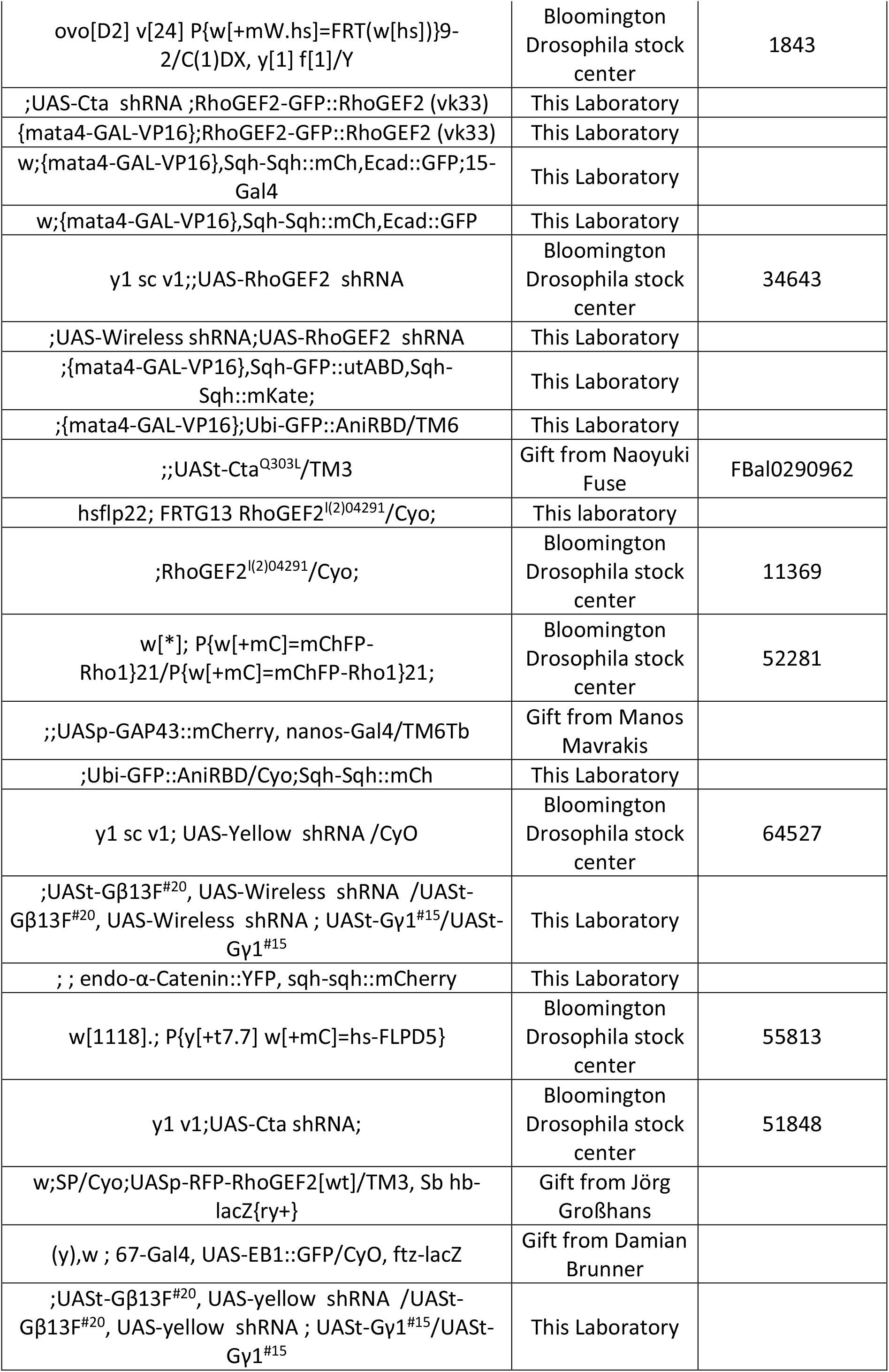
Drosophila strains used in this study

**Supplementary table 2.**
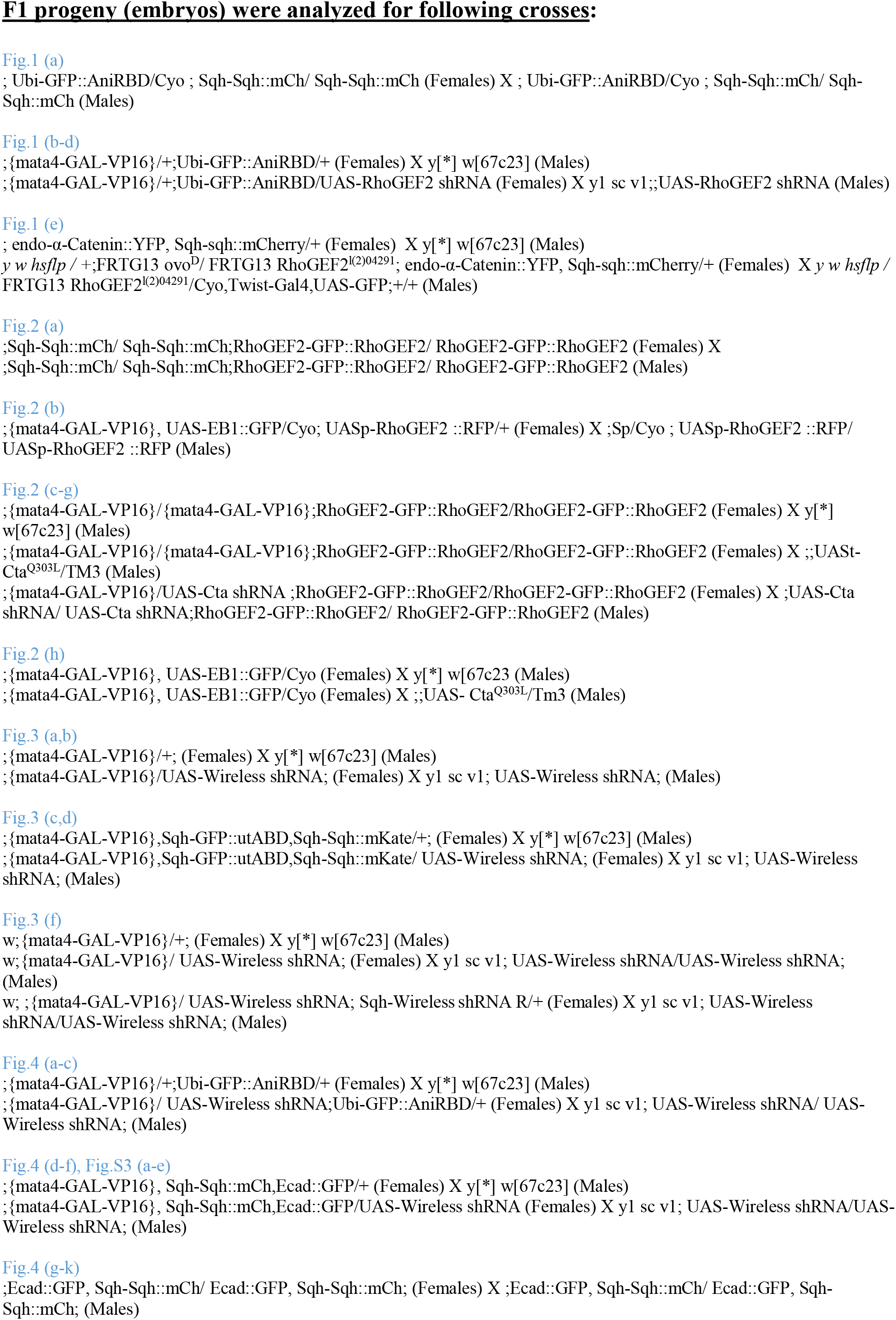
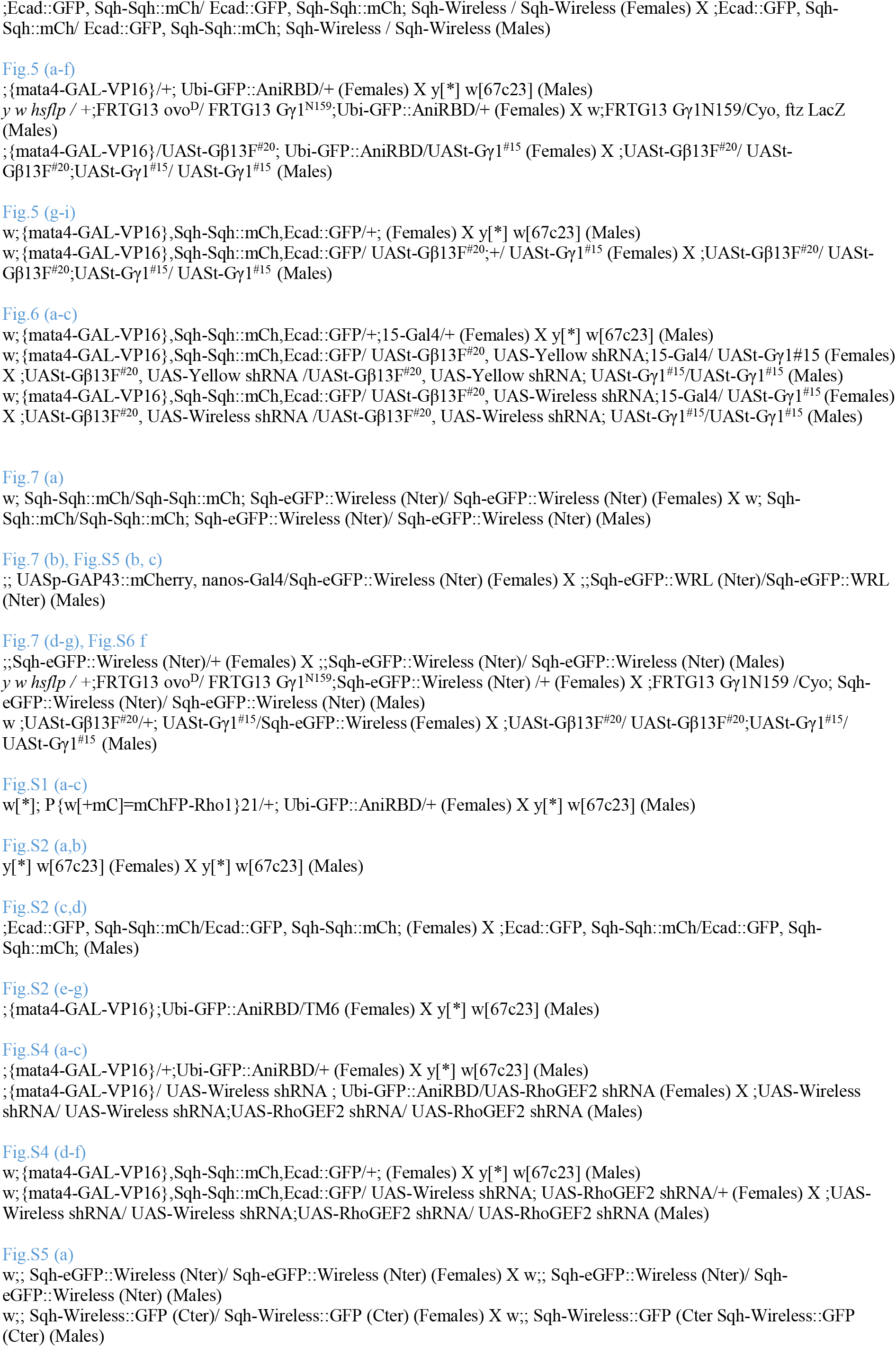
Fly crosses performed in this study

